# SUMO4 promotes SUMO deconjugation required for DNA double-strand break repair

**DOI:** 10.1101/2022.03.23.485504

**Authors:** Alexander J. Garvin, Alexander J. Lanz, Katarzyna Starowicz, George E Ronson, Matthew J.W. Mackintosh, Alexandra K. Walker, Yara Aghabi, Hannah MacKay, Ruth M. Densham, Aneika C. Leney, Joanna R. Morris

**Affiliations:** Birmingham Centre for Genome Biology and Institute of Cancer and Genomic Sciences, College of Medical and Dental Sciences, University of Birmingham, B15 2TT, UK; SUMO Biology Lab, School of Molecular and Cellular Biology, Faculty of Biological Sciences, University of Leeds, Leeds, LS2 9JT; School of Biosciences, College of Life and Environmental Sciences, University of Birmingham, B15 2TT, UK

**Keywords:** SUMO, SUMO4, DNA repair, Homologous Recombination, RAP80, SENP1

## Abstract

The amplitudes of small-modifier protein signalling through ubiquitin and the Small Ubiquitin-like Modifiers, SUMO1-3, are critical to the correct phasing of DNA repair protein accumulation, activity, and clearance, and for the completion of mammalian DNA double-strand break (DSB) repair. However, how SUMO-conjugate signalling in the response is delineated is poorly understood. At the same time, the role of the non-conjugated SUMO protein, SUMO4, has remained enigmatic. Here we reveal that SUMO4 is required to prevent excessive DNA-damage-induced SUMOylation and deleterious over-accumulation of RAP80. Mechanistically we show SUMO4 acts independently of its conjugation and potentiates SENP1 catalytic activity. These data identify SUMO4 as a SUMO deconjugation component and show SUMO4:SENP1 are critical regulators of DNA-damage-induced SUMO signalling.

**Highlights:**

- The roles of the SUMO family members, SUMO1-4, in DSB repair are not redundant.
- SUMO4 conjugation is deleterious.
- SUMO4 promotes SENP1 catalytic activity.
- SUMO4:SENP1 restrict SUMO-signalling and accumulation of the BRCA1-A complex.

## Introduction

DNA double-strand-breaks (DSBs) are severely toxic genomic lesions. Our cells have evolved the means to signal DNA breaks in order to launch appropriate pathways to repair these lesions. DSBs induce histone modifications and a recruitment cascade of repair proteins regulated by histone and repair protein modifications of phosphorylation, acetylation, methylation, ubiquitination and the conjugation of Small Ubiquitin-like Modifiers, SUMO1-3 (SUMOylation). The sequencing, duration and amplitude of post-translational modification signals in the response are critical to directing both a favourable environment for repair and the correct recruitment of repair proteins.

SUMO protein paralogs include SUMO2 and SUMO3, which share 95% identity (hereafter SUMO2/3), and SUMO1, which shares 48% and 46% identity with SUMO2 and SUMO3, respectively ^1^. A fourth SUMO protein, SUMO4 has 86% homology to SUMO2, and antibodies that detect SUMO4 also cross-react with SUMO2/3 across a range of detection formats ^2^. The *SUMO4* gene is a retrogene, its mRNA has been identified by ribosome-profiling across many cell types, chiefly lymphoblastoid and fibroblast cell lines at folds lower than *SUMO2* or *SUMO3* mRNAs ^3^. Despite its low abundance, proteomics studies have identified amino acid sequences unique to SUMO4, including “A**N**EKP**T**E**E**VKTENN**N**HINLK”, and “TENN**N**HINLK” (SUMO4-specific amino acids in bold), confirming its presence *in vivo* ^4–6^. Inactive precursors of SUMOs are matured by SENtrin-specific proteases (SENPs) to expose the di-glycine C-terminal motif required for conjugation ^7^. SUMO4 contains a C-terminal proline reported to suppress maturation ^8^. Some reports indicate SUMO4 is not conjugated ^8^, while others suggest conjugation ^9–11^ and the possibility that it is matured by non-SENP enzymes ^9^, so the relevance of SUMO4 to cellular biology is currently enigmatic. Conjugation of SUMOs to the ε-amino groups of target lysines commonly uses a three-enzyme cascade (E1-E2-E3) analogous to the enzyme architecture for ubiquitin modification ^12^. For many substrates, SUMOylation is highly dynamic. Conjugate liability is due in part to the rapid isopeptidase activity of SENP1-3 and SENP5-7 proteins ^13^. A small number of non-SENP deSUMOylating enzymes have also been identified ^14,15^.

The balance between SUMOylation and deSUMOylation is altered by exposure to environmental or metabolic stresses, including DSBs ^16–20^. In *S. cerevisiae* SUMOylation of homologous recombination repair factors occurs through a DNA-bound SUMO E3 ligase ‘spray’, and in mammalian cells, SUMOylation of protein complexes is consistent with protein group modification ^21–23^. The mammalian SUMO E3 ligases PIAS1, PIAS4 and CBX4 drive SUMOylation induced by DNA breaks ^17,24–26^, with some indications that SUMO1 conjugation predominates early in the DSB signalling cascade, and SUMO2/3 later ^24,26,27^. SUMOylation, in large part, acts by increasing interactions between SUMOylated factors and the SUMO-interacting domains (SIMs and other interfaces ^28–31^) of partner proteins, which can include SUMO-targeting ubiquitin ligases (StUbLs), leading to extraction and degradation of modified proteins ^23,32–35^. Thus deSUMOylation, for example of MDC1, restricts StUbL access and regulates the duration of MDC1 localization and, in turn, DSB signalling ^27^. Consistent with the proposed role of SUMO as a “molecular glue”^21–23^, SUMOylation at DSBs also drives the recruitment of various proteins, including the BRCA1-A complex ^36,37^; the deubiquitinating enzyme, Ataxin-3^38^; and the nuclease scaffold SLX4 ^39,40^ and promotes interactions between RPA-RAD51-BRCA2^41,42^ and XRCC4 with regulators of non-homologous end-joining ^43,44^.

Here, we compare the influence of each SUMO protein on DSB signalling, finding that each, surprisingly including SUMO4, has distinct roles in the DSB response. We investigate the role of SUMO4 and find that its conjugation is incompatible with its function in DSB repair. We find that SUMO4 promotes SENP1 deSUMOylase activity and, consequently, is required to limit SUMOylation induced following DNA damage. Further, our data indicate that SUMO4:SENP1 has no influence on SUMOylation/deSUMOylation regulating MDC1 but acts to suppress SUMOylation directed by PIAS1 responsible for RAP80 recruitment, providing further evidence of separable phases of SUMO DSB signalling in mammalian cells. SUMO4 is consequently required for DNA repair, genome stability and responses to genotoxic stress. This study identifies a surprising, non-canonical role for SUMO4 as a novel deconjugation component of the SUMO system. It also shows that SUMO4 is critical to correct SUMO-signalling amplitude following DSB generation and for the promotion of DNA repair.

## Results

### SUMO proteins are non-redundant in DSB repair, and SUMO4 plays a distinct role

To compare the roles of each SUMO protein, we generated siRNA sequences specifically targeting each, including SUMO4, over the similar SUMO2/3 (Figure 1A and S1A). Many antibodies raised to SUMO4 cross-react with SUMO2 and SUMO3, and most of the signal discerned by these antibodies is due to SUMO2/3^2^. We were able to precipitate exogenous SUMO4 slightly better than exogenous SUMO3 when both proteins were highly expressed using one of these antibodies, IOO-19 (Figure S1B), and liquid chromatography-mass spectrometry of immunoprecipitated endogenous proteins revealed SUMO4-specific peptides (Figure S1C-E). These findings are consistent with low-level SUMO4 protein expression previously reported ^4–6^. A SUMO4 siRNA-sensitive band was discernible by western blot (Figure S1F), and expression of exogenous 6xHis-HA-SUMO4 in U2OS cells was suppressed by SUMO4 siRNA (Figure S1G). These data indicate SUMO4 siRNA suppresses SUMO4 protein expression. *SUMO4* is located within the final intron of the *TAB2* (*MAP3K7IP2*) gene and TAB2 itself has a reported signalling role in DSB repair ^45^. Importantly we noted that SUMO4 siRNAs had little impact on TAB2 protein expression (Figure S1H), suggesting siRNAs directed against SUMO4 are unlikely to be suppressing TAB2. Next, we compared the effects of different SUMO siRNAs on cell proliferation and cycle distribution of U2OS cells. In contrast to SUMO1 or SUMO2 siRNA treatment, SUMO4 siRNA did not result in a significant decrease in cell number (Figure S1I), and SUMO4 siRNA slightly increased the proportion of cells in S-phase over G1 (Figure S1J). We tested DNA repair outcomes of integrated reporters for homologous recombination (gene conversion) and non-homologous end-joining, noting that all SUMO siRNAs, except that to SUMO3, reduced repair outcomes (Figure 1B). SUMO1-4 siRNA treatment also delayed clearance of phosphorylated serine-139 H2AX (γH2AX) foci following exposure to ionizing irradiation (IR) in both EdU negative and positive (S phase) U2OS cells (Figure 1C and S1K).

**Figure 1.**
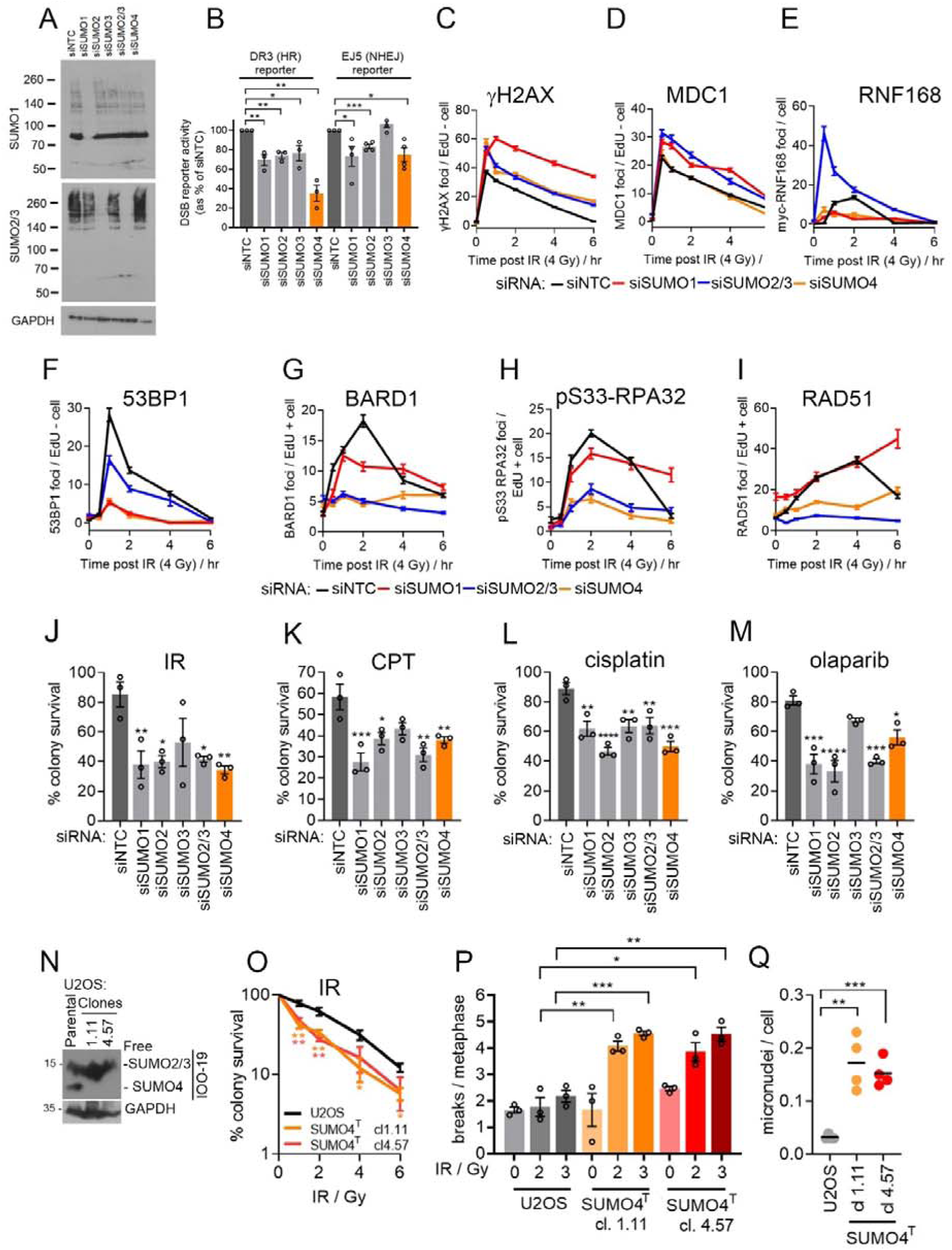
SUMO4 is required for DSB repair. **A)** Immunoblot of lysates from U2OS treated with indicated siRNAs, NTC: non-targeting / luciferase or to SUMO isoforms. SUMO1-4 are targeted by two siRNA sequences each, siSUMO2/3 uses a combination of SUMO2 and SUMO3 sequences. The SUMO2/3 antibody detects both SUMO2 and SUMO3. **B)** U2OS DR3, Homologous recombination (HR) reporter or EJ5 non-homologous end-joining (NHEJ) reporter cells were co-transfected with siRNA and DNA for the restriction enzyme *SceI,* and RFP as a transfection marker. Events were gated on RFP (*SceI* transfection) and GFP (repair reporter). The repair efficiency was calculated relative to siNTC/RFP transfected cells expressed as 100%. N=3 (HR) N=4 (NHEJ). Statistical significance was assessed by two-tailed *t*-test between siNTC and siSUMO. **C)** Number of γH2AX foci in U2OS cells treated with indicated siRNA for 48 hr before (4 Gy) and fixed at the indicated times. The were counted in EdU negative and positive cells (Supplemental Figure 1b). N=∼150 cells from a total of 3 experimental repeats per condition. **D)** Quantification of MDC1 foci **E)** myc-RNF168 foci (in U2OS expressing doxycycline inducible (48 hr) myc-RNF168), **F**) 53BP1, **G**) BARD1 foci, **H**) RPA32 phospho-S33 foci and **I**) RAD51 foci in U2OS cells treated with indicated siRNA for 48 hr before (4 Gy) and fixed at the indicated times. Foci kinetics for 53BP1 in EdU + cells are shown Supplemental Figure 1c. **J-M)** Cell survival of U2OS measured by colony assay following treatment with th indicated siRNAs for 48 hours before treatment **J**) 2 Gy IR, **K**) 1 μM camptothecin (CPT), **L**) 1 μM cisplatin, **M**) 10 μM olaparib. All drug treatments were for 2 hrs. Statistical significance was assessed by two-tailed *t*-test between siNTC and siSUMO. **N)** Immunoblot of lysates from parental and SUMO4 edited U2OS clones, cl.1.11 (gRNA #1) and cl.4.57 (gRNA #4) probed with anti-SUMO2/3/4 monoclonal IOO-19 or GAPDH. **O)** Cell survival of U2OS measured by colony assay of parental U2OS^FlpIn^ and SUMO4^T^ cl.1.11 (gRNA #1) and cl.4.57 (gRNA #4) treated with IR at the indicated doses. * denote statistical difference for each dose between parental U2OS and SUMO4^T^ clones, determined by two-tailed t-test. **P)** Number of breaks (chromatid and chromosome) per metaphase from three independent experiments in untreated or in cells analyzed 24 hrs after exposure to 2 or 3 Gy IR. Statistical significance was assessed by two-tailed *t*-test between the parental U2OS and SUMO4^T^ for each IR dosage. **Q)** Mean micronuclei number per cell from 4 independent repeats of IR treated (4 Gy, fixed 6 hr later) parental U2OS^FlpIn^ and SUMO4^T^ clones 1.11 and 4.57. ∼ 100 nuclei per condition, 4 experimental repeats.

Next, we assayed DSB repair protein foci kinetics in SUMO-depleted U2OS cells treated with IR, first assessing MDC1, a repair factor that requires SUMOylation for clearance from DSBs in G1^27,32–34,38^. MDC1 foci resolution was delayed following the depletion of SUMO1 or SUMO2/3 but was unaffected by SUMO4 siRNA (Figure 1D). In contrast, the foci formation of the ubiquitin E3 ligase RNF168 was suppressed in SUMO1 and SUMO4-depleted cells, and SUMO2/3-depletion resulted in excessive RNF168 accrual, particularly at early time points (Figure 1E). When we assessed the recruitment of two downstream readers of the RNF168-generated N-terminally ubiquitinated H2A, 53BP1 and BARD1 ^46–48^, we found that both were reduced after SUMO4 depletion (Figures 1F-G and S1L). We then tested RPA32 pSer33 foci as an indication of DNA end-resection ^49^, and RAD51 foci as an indication of the generation of homologous recombination intermediates. Treatment with SUMO4 and SUMO2/3 siRNA reduced both markers following IR (Figure 1H-I). In the assessment of the responses to genotoxic agents, depletion of each SUMO isoform (with the exception of SUMO3, after IR or olaparib) increased U2OS cell sensitivity to IR, camptothecin (CPT), cisplatin, and the PARP1/2 inhibitor, olaparib (Figure 1J-M). These data confirm the non-redundant roles of SUMO1 and SUMO2/3 proteins in the DNA damage response and show that SUMO4 targeting impacts DSB signalling in a manner distinct from the depletion of SUMO1 or SUMO2/3.

To explore the role of SUMO4 further, we tested SUMO4 siRNA treatment of HeLa and NCI-H1299 cells and found that it also sensitized these cells to IR, CPT and cisplatin (Figure S1M & N), suggesting the requirement for SUMO4 is not restricted to U2OS cells. We next disrupted the *SUMO4* locus using two different gRNAs in independent U2OS clones (Figures S1O), creating stop-codon inducing edits part-way through the coding region of *SUMO4;* after codon 45 on both alleles of clone 1.11 and after codon 72 (allele A) and 56 (allele B) of clone 4.57. Western analysis of lysates from these clones indicated that the SUMO4 siRNA-sensitive band was absent (Figure 1N). No SUMO4-specific peptides were identified from clone 4.57, whereas the N-terminal peptide “PTEEVKTENNNHINLK” was identified in clone 1.11 (Figure S1C). Genes lacking the intron structure to regulate the decay of transcripts with premature stop codons often retain mutant mRNA expression^50^, and thus an expression of a severely truncated SUMO4 protein is expected, these clones are called U2OS-SUMO4^T^ (truncated) hereafter. To test whether cells bearing *SUMO4* disruption exhibited similar phenotypes to those treated with SUMO siRNA, we assessed sensitivity to DNA-damaging agents and the recruitment of proteins after IR-exposure. Both genomically targeted clones were sensitive to treatments with IR, CPT, and cisplatin (Figures 1O and S1P) and these cells also reproduced defects in 53BP1/RAD51 foci number after IR-exposure (Figure S1Q). Importantly, the treatment of these cells with SUMO4 siRNA produced no additional defects in DSB signalling or survival (Figure S1Q & R), indicating *SUMO4* gene edits and SUMO4 siRNA disrupt the same cellular feature. As for siRNA treatment, the *SUMO4*-disrupted clones displayed normal levels of TAB2 protein (Figure S1S). The genetic, immunoblot and functional data together indicate that the edited clones lack full SUMO4 activity.

Intriguingly, we noted that SUMO4^T^ cells retained a low level of IR-induced 53BP1 and RAD51 foci, which could be further reduced by RNF8/RNF168 or BRCA2 siRNA, respectively, suggesting that DSB repair signalling in these cells is substantially reduced, rather than absent (Figure S1T). To assess whether SUMO4 suppresses the chromosomal consequences of DNA damage, we analyzed metaphase spreads and noted increased chromosomal breaks in IR-treated SUMO4^T^ cells *versus* the parental U2OS (Figure 1O). Additionally, micronuclei, a marker of fragmented chromosomes, were increased in both IR-treated SUMO4^T^ and siSUMO4 cells (Figures 1P and S1U). In summary, SUMO4 has little influence on MDC1 kinetics but promotes RNF168, 53BP1 and RAD51 foci accumulation. Consistent with these observations, SUMO4 promotes measures of HR and NHEJ, is required for cellular resistance to genotoxins, and the suppression of genomic instability.

### SUMO4 function is suppressed by its conjugation

To address whether conjugation proficiency relates to SUMO4 function in DSB repair, we tested a series of siRNA-resistant SUMO4 mutants. These were in two categories: those designed to ensure conjugation incompetence and those altered to enable the mutant protein to be attached to target lysines through the SUMO conjugation pathway (conjugation proficient - Figure 2A). The conjugation incompetent mutants were those bearing a stop codon before the glycine residues that are essential for conjugation T91X (TX) or in which the glycine residues were mutated to alanines, G92A/G93A (GA). Proline 90 within the SUMO4 C-terminal tail is proposed to interfere with SENP-mediated exposure of the di-glycine motif, suppressing SUMO4 maturation ^8^. We generated conjugation-proficient mutants through SUMO4-P90Q (PQ) mutation, reflecting the Gln at this position in SUMO2, and made an artificially matured variant bearing a stop codon after the glycine residues V94X (VX). As a further control, we combined the V94X and G92/G93A mutations (GA/VX), expected to be incompetent for lysine conjugation (Figure 2A). As anticipated, only the SUMO4-PQ and SUMO4-VX mutants formed high molecular weight smears, indicating conjugation by the cellular SUMOylation machinery, whereas SUMO4-GA, SUMO4-TX and SUMO4-GA/VX migrated as a monomer similar to SUMO4-WT (Figure 2B) suggesting both WT SUMO4 and these mutants are not conjugated. Further, in examining the subcellular localization of the mutant proteins, we found that conjugation-proficient mutants were nuclear, whereas conjugation-deficient forms showed cytoplasmic and nuclear localization resembling the SUMO4-WT protein (Figure S2A).

**Figure 2.**
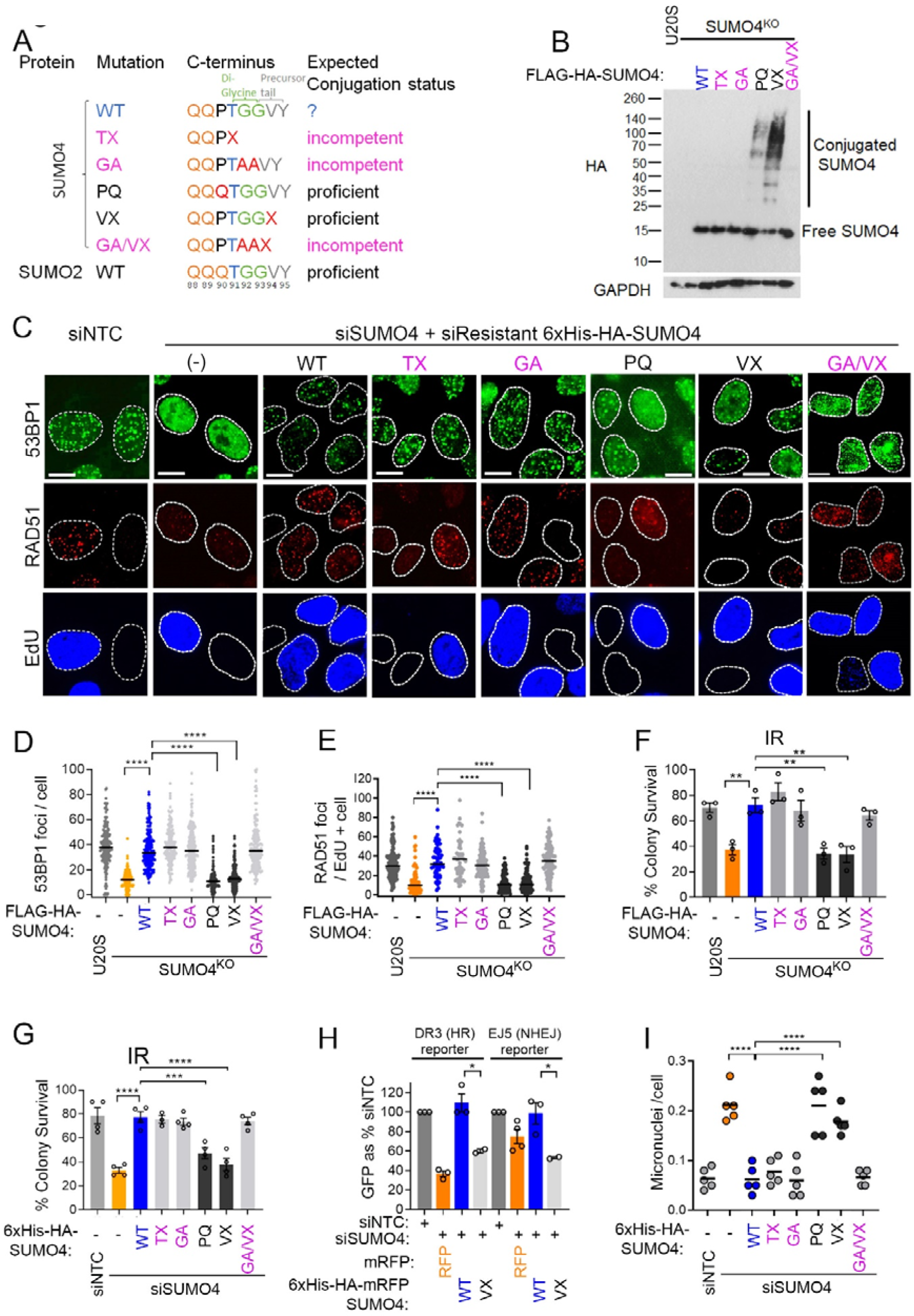
Conjugation independent role of SUMO4 in DSB repair. **A)** Illustration of the C-terminal tail of WT SUMO4 and the generated mutants. The di-Glycine repeat required for conjugation in SUMO1-3 is highlighted in green. The precursor tail removed by SENPs during Pro-SUMO maturation is highlighted in grey. Stop codons are shown as X. Numbering relates to SUMO4 sequence and the SUMO2 C-terminus is shown for comparison. The expected conjugation proficiency is illustrated **B)** Retardation of some SUMO4 mutants expressed in SUMO4^T^ cl1.11 cells by induction with doxycycline for 48 hrs to induce FLAG-HA SUMO4 protein expression. Lysates were immunoblotted with HA antibody and GAPDH loading control. **C)** Representative images of U2OS cells treated with SUMO4 siRNA and doxycycline for 48 hrs to induce expression of siRNA-resistant 6xHis-HA-SUMO4 variants before IR treatment (4 Gy), fixed 2 hr later and immunostained for 53BP1 or RAD51. Cells were pulsed with 10 μM EdU 30 min prior to fixation to label S phase cells. Scale bar = 10 μm. **D-E)** Quantification of 53BP1 (**D**) or RAD51 (**E**) foci in SUMO4^T^ cl1.11 cells doxycycline-treated for 48 hr to induce FLAG-HA SUMO4 expression before IR treatment (4 Gy) and fixed 2 hr later for immunostaining. N=∼150 cells per condition from 3 experiments. RAD51 foci are quantified in EdU + cells. **F)** Cell survival of SUMO4^T^ cl1.11 treated with 48 hr doxycycline to induce FLAG-HA-SUMO4 proteins before irradiation (2 Gy) measured by colony assay. N=3. Statistical differences determined by one way ANOVA. **G)** Cell survival of U2OS treated with SUMO4 siRNA for 48 hr with concurrent doxycycline treatment to induce expression of siRNA resistant 6xHis-HA SUMO4 before exposure to 2 Gy IR, measured by colony assay. Statistical differences determined by one way ANOVA. **H)** DNA repair out comes in U2OS DR3 (HR reporter) or EJ5 (NHEJ reporter) cells co-transfected with SUMO4 siRNA and DNA for *SceI* and mRFP or His-HA-mRFP-SUMO4. Events gated on GFP (repair reporter) and RFP (positive transfection). Repair efficiency is calculated relative to siNTC/RFP transfected cells expressed as 100%. Statistical differences were determined by two-tailed *t*-test. **I)** Micronuclei number per cell in U2OS treated with SUMO4 siRNA for 48 hr with concurrent doxycycline treatment to induce expression of siRNA resistant 6xHis-HA-SUMO4 proteins. N=5. Statistical differences determined by one way ANOVA.

We tested each mutant in complementation assays in SUMO4^T^ cells or U2OS treated with SUMO4 siRNA, using 53BP1 and RAD51 as markers of DSB signalling. Conjugation-incompetent SUMO4 variants (GA and TX) restored 53BP1 and RAD51 foci to control/WT levels in both backgrounds. In contrast, the conjugation-proficient mutants (PQ and VX) did not rescue the reduced 53BP1 or RAD51 foci in SUMO4-deficient cells. Further, converting the conjugation-proficient SUMO4-VX to a conjugation-incompetent form by mutation of the di-Gly (GA/VX) restored DSB signalling (Figures 2C-E and S2B-C). Colony survival analysis of complemented cells showed that the conjugation-incompetent (WT, TX, GA and GA/VX), but not the conjugation-proficient mutants (PQ and VX), were able to restore resistance to IR, CPT, cisplatin and olaparib (Figures 2F-G and S2D-I). Similarly, the conjugation-proficient mutant, SUMO4-VX, was unable to restore GFP expression in either HR or NHEJ *SceI* reporter assays (Figure 2H). Suppression of micronuclei formation mirrored the SUMO4 mutants’ ability to promote DSB signalling and survival (Figure 2I). To address whether the impact of conjugation-proficient mutants reflects a deleterious gain of function(s), we examined their over-expression. Whereas overexpression of SUMO4-WT in parental U2OS had no impact on cell survival, the expression of conjugation-proficient SUMO4-PQ and SUMO4-VX variants reduced cell survival after exposure to IR (Figure S2J), suggesting that SUMO4ylation is deleterious to the normal response to DNA damage. We next tested whether the expression of a conjugation-defective form of an alternative SUMO, SUMO2, can complement SUMO4^T^ cells. Indeed, expression of conjugation deficient SUMO2-GA (G92A/G93A) in SUMO4^T^ cells improved 53BP1/RAD51 foci and cell survival after exposure to IR (Figure S2K-M), suggesting either SUMO4 performs a role shared by free SUMO2 or that increased free SUMO2 suppresses the harmful consequences of absent SUMO4. Collectively, these results show that the conjugation of SUMO4 is incompatible with its function in DSB repair and indicate that non-physiological SUMO4 conjugates reduce survival after damage.

### SUMO4 requires its SIM binding patch for DSB repair activity

SUMO4 shares several features with its SUMO2 ancestor, including the surface that proteins with SUMO interacting motifs (SIMs) interact with (Figure 3A). To determine if this feature is relevant to the DSB repair-promoting function of SUMO4, we generated mutations analogous to the previously characterized SUMO2:SIM interaction mutant ^51^; SUMO4-Q31A/F32A/I34A (QFI-A) and found that while SUMO4-WT protein was pulled down by SIM-bearing peptides (derived from PIAS1 and SLX4 proteins), SUMO4-QFI-A was absent in the SIM peptide pull-down (Figure S3A). We found SUMO4-WT and SUMO4-QFI-A showed similar subcellular localization (Figure S3B). However, SUMO4^T^ or SUMO4 siRNA treated cells complemented with SUMO4-QFI-A were defective in 53BP1 and RAD51 foci formation after IR treatment (Figures 3B-E and S3C-D), were sensitive to DSB inducing agents in colony survival assays (Figures 3F-G and S3E-F) and showed poor HR repair in integrated GFP gene-conversion assays (Figure 3H). Micronuclei formation was also not suppressed in SUMO4-QFI-A complemented SUMO4 siRNA-treated or SUMO4^T^ cells (Figure 3I). These data implicate the SIM-SUMO4 interaction in the repair of DNA breaks.

**Figure 3.**
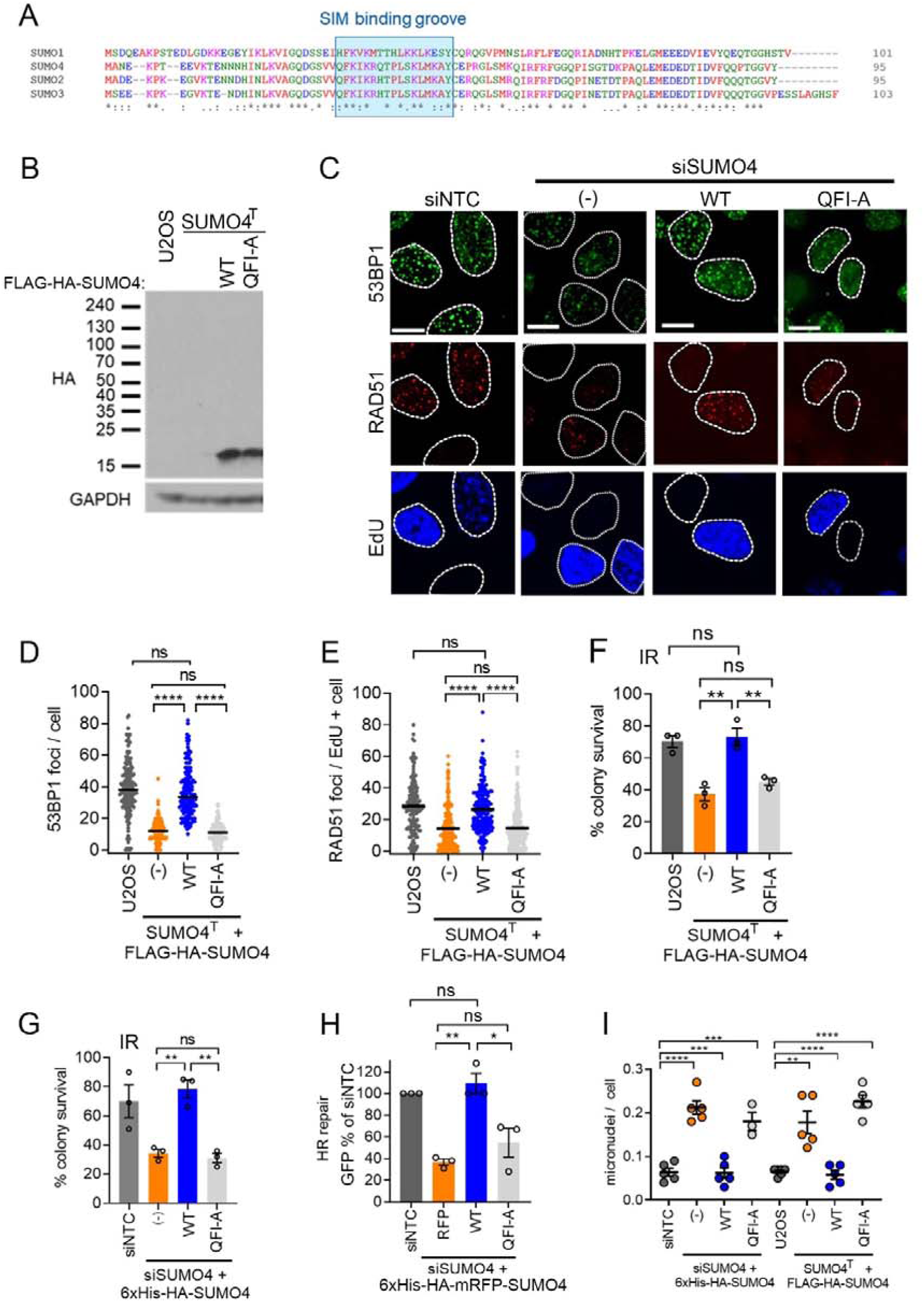
SUMO4 requires its SIM binding groove to promote DSB repair activity. **A)** Alignment using Clustal Omega of human SUMO1 (P63165), SUMO2 (P61956), SUMO3 (P55854) and SUMO4 (Q6EEV6). The SIM binding groove is highlighted in blue. **B)** Immunoblot of SUMO4^T^ cl1.11 cells expressing FLAG-HA-SUMO4-WT or FLAG-HA-SUMO4-QFI-A mutant, probed with HA-antibody and GAPDH loading control. **C)** Representative images of 53BP1 and RAD51 foci in U2OS cells treated with siSUMO4 and and doxycycline for 48 hrs to induce expression of siRNA-resistant 6xHis-HA-SUMO4 variants before IR treatment (4 Gy) fixed 2 hrs later and stained. EdU marks S phase cells. Scale bars = 10 μm. **D)** U2OS SUMO4^T^ cl1.11 complemented with FLAG-HA-SUMO4-WT or FLAG-HA-SUMO4-QFI-A were treated with dox for 48 hr prior to IR (4 Gy) and fixed 2 hr later. The parental U2OS^FlpIn^ were used as controls. N= a total of ∼ 150 cells per condition from 3 experimental repeats. Statistical differences were determined by two-tailed *t*-test. **E)** RAD51 foci were scored in EdU-positive cells treated as in d). Statistical differences were determined by two-tailed *t*-test. **F)** Cell survival of U2OS SUMO4^T^ cl1.11 treated with doxyclycine for 48 hr prior to IR (2 Gy) to induce with FLAG-HA-SUMO4-WT or FLAG-HA-SUMO4-QFI-A before plating for colony growth. Statistical differences were determined by two-tailed *t*-test. **G)** Cell survival of U2OS cells treated with control (NTC) siRNA or siRNA to SUMO4 and doxycycline-treated to express 6xHis-HA-SUMO4 proteins before IR (2 Gy) and plating for colony assay. Statistical differences were determined by two-tailed *t*-test. **H)** U2OS-DR3 cells siRNA depleted of SUMO4 and transfected with *SceI*, RFP or 6xHis-HA-mRFP-SUMO4 simultaneously. Cells were gated on GFP/RFP positivity, and the % GFP-positive cells are shown relative to those treated with siNTC. N=3 experimental repeats performed in triplicate. Statistical differences were determined by two-tailed *t*-test. **I)** Quantification of micronuclei number per cell in 6xHis-HA-SUMO4 complemented siSUMO4 U2OS and or FLAG-HA SUMO4 complemented SUMO4^T^ cl1.11 cells. N=5 with ∼ 100 nuclei counted per experiment. Statistical differences were determined by two-tailed *t*-test.

### SUMO4 maintains SUMO1-3 homeostasis

We next assessed the impact of SUMO4 loss on total cellular SUMOylation and on PML nuclear bodies (PML-NBs), which are cellular features sensitive to altered SUMO conjugation ^52,53^. In SUMO4^T^ cells, we observed increased SUMO1 and SUMO2/3 conjugates in untreated and IR-treated cells (Figures 4A and S4A). In cells treated with SUMO4 siRNA, the nuclear immunofluorescence intensity of SUMO2/3 was elevated in untreated and IR-treated cells and SUMO1 intensity was elevated following IR-treatment (Figure 4B & C). Complementation with SUMO4-WT suppressed these measures, but the expression of the SUMO4-QFI-A or conjugation-proficient mutants did not (Figure 4D-E and S4B-C). We also noted an increase in both the number of PML-NBs, and in bodies containing the PML-NB component, Sp100, in cells lacking SUMO4, which could be restored to control levels by expression of SUMO4-WT, but not SUMO4-QFI-A (Figure S4D-G).

**Figure 4.**
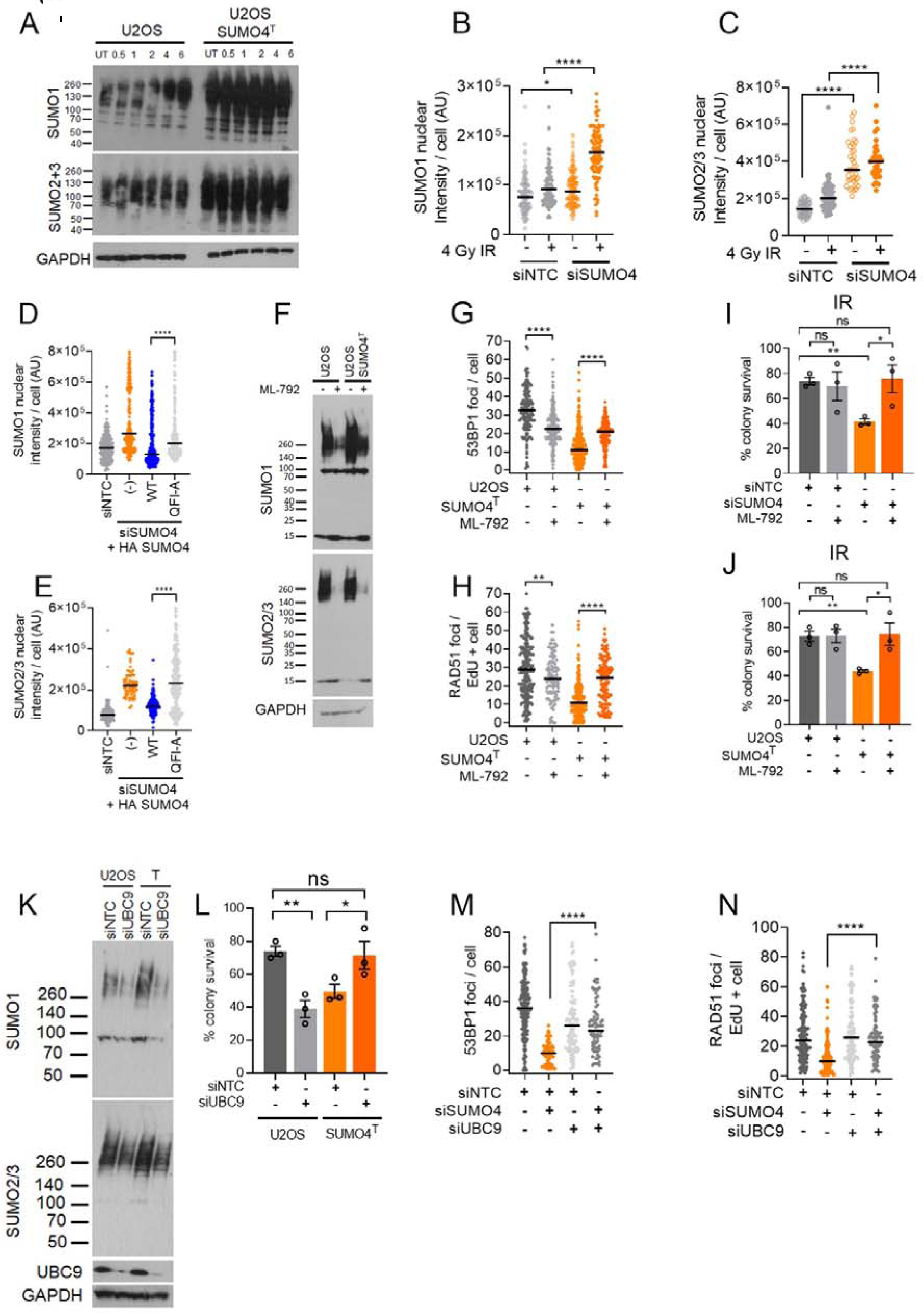
SUMO4 maintains SUMO1-3 homeostasis. **A)** High molecular weight SUMO in whole cell lysates from U2OS or U2OS SUMO4^T^ cl1.11 treated with 4 Gy IR, lysed at indicated time points, immunoblotted for SUMO1, SUMO2/3 and GAPDH loading control. **B and C)** Quantification of indirect immunofluorescence intensity of SUMO1 (**B**) and SUMO2/3 (**C**) in U2OS treated with non-targeting control siRNA (siNTC) or SUMO4 siRNA (siSUMO4) for 48 hr followed by 4 Gy IR, 2 hrs later cells were pre-extracted to remove soluble material and fixed before immunostaining with the relevant antibodies. Fluorescence intensity per cell is shown in arbitrary units (AU) from ∼100 cells per condition. Statistical differences were determined by two-tailed *t*-test. **D and E)** Quantification of indirect immunofluorescence intensity of SUMO1 (**D**) or SUMO2/3 (**E**) in U2OS treated with SUMO4 siRNA for 48 hr and doxycycline for 16 hr, or not (-), to induce 6xHis-HA-SUMO4-WT or 6xHis-HA-SUMO4-QFI-A expression, followed by IR (4 Gy) treatment and fixation at 2 hr. Cells were extracted, fixed and stained as for B) and C). N= ∼150 cells per condition from a total of three experiments. Statistical differences were determined by two-tailed *t*-test. **F)** Assessment of high molecular weight SUMO from lysates of U2OS or U2OS SUMO4^T^ cl1.11 cells treated with 1 μM ML-792 (+) or DMSO (-) by immunoblot with antibodies to SUMO1, SUMO2/3. UCB9 and GAPDH immunoblots are also shown. **G and H)** Quantification of 53BP1 foci (**G**) and RAD51 foci (**H**) in U2OS SUMO4^T^ cl1.11 treated with 1 μM ML-792 or DMSO for 1 hour before irradiation (4 Gy), fixed 2 hr later, and immunostained for the relevant proteins. n = ∼150 cells per condition from 3 experiments. Statistical differences were determined by two-tailed *t*-test. **I)** Cell survival measured by colony assay of U2OS treated with siNTC or siSUMO4 and either 1 μM ML-792 or DMSO for 1 hour before irradiation (2 Gy). n =4. Statistical differences were determined by two-tailed *t*-test. **J)** Cell survival measured by colony assay of U2OS SUMO4^T^ cl1.11 or parental U2OS treated with 1 μM ML-792 or DMSO for 1 hr before irradiation (2 Gy) n =4. Statistical differences were determined by two-tailed *t*-test. **K)** Assessment of high molecular weight SUMO from lysates of U2OS or U2OS SUMO4^T^ cl1.11 cells treated with siNTC or UBC9 siRNA (siUBC9) by immunoblot with antibodies to SUMO1, SUMO2/3. UCB9 and GAPDH immunoblots are also shown. **L)** Cell survival measured by colony assay of U2OS or SUMO4^T^ cl1.11 cells treated with UBC9 siRNA for 48 hr prior to IR (2 Gy) and plating. N=3. Statistical significance was assessed by one way ANOVA. **M and N)** Quantification of 53BP1 foci (**M**) and RAD51 foci (**N**) in U2OS cells treated with siNTC or siSUMO4 with or without UBC9 siRNA (siUBC9) for 48 hr before irradiation (4 Gy). Cells were fixed after 2 hrs and stained for the detection of 53BP1 or RAD51. N= >75 cells per condition from a total of three experiments. RAD51 foci were scored in EdU + cells. Statistical differences were determined by two-tailed *t*-test.

To test whether the increased SUMOylation observed following SUMO4 loss influences the poor DSB repair proficiency of SUMO4-deficient cells, we next manipulated SUMO conjugation. We first tested short-term treatment with the SUMO E1 inhibitor ML-792 ^54^ (Figure 4F). Remarkably ML-792 treatment improved 53BP1 and RAD51 foci accrual and resistance to IR in both SUMO4 siRNA-treated and SUMO4^T^ cells (Figure 4G-J). To challenge this idea further, we partially depleted the SUMO E2, UBC9. This treatment also improved IR resistance in SUMO4^T^ cells (Figure 4K-L) and increased the accumulation of 53BP1 and RAD51 foci in SUMO4 siRNA-treated cells (Figure 4M & N). Analysis of cells treated with siRNAs targeting PIAS1-4 SUMO E3 ligases determined that PIAS1 siRNA treatment improved 53BP1 and RAD51 foci numbers and resistance to IR, CPT and olaparib in SUMO4^T^ cells (Figure S4H-M). These data suggest hyperSUMOylation underlies defective DSB signalling in the absence of SUMO4.

### SUMO4 promotes SENP1 protease activity

To determine the cause of the increased SUMO conjugates in SUMO4-deficient cells, we prepared cellular extracts without the cysteine protease inhibitors that are usually included to suppress the protease-mediated cleavage of Ub/Ubl conjugates. We examined the loss of high-molecular-weight SUMO1 and SUMO2/3 with time. SUMO4^T^ cells showed a slower rate of reduction of high molecular weight SUMO-conjugates, which could be restored by the expression of exogenous SUMO4-WT (Figure 5A-D), suggesting a reduced ability to deconjugate SUMO1-3 in SUMO4^T^ extracts.

**Figure 5.**
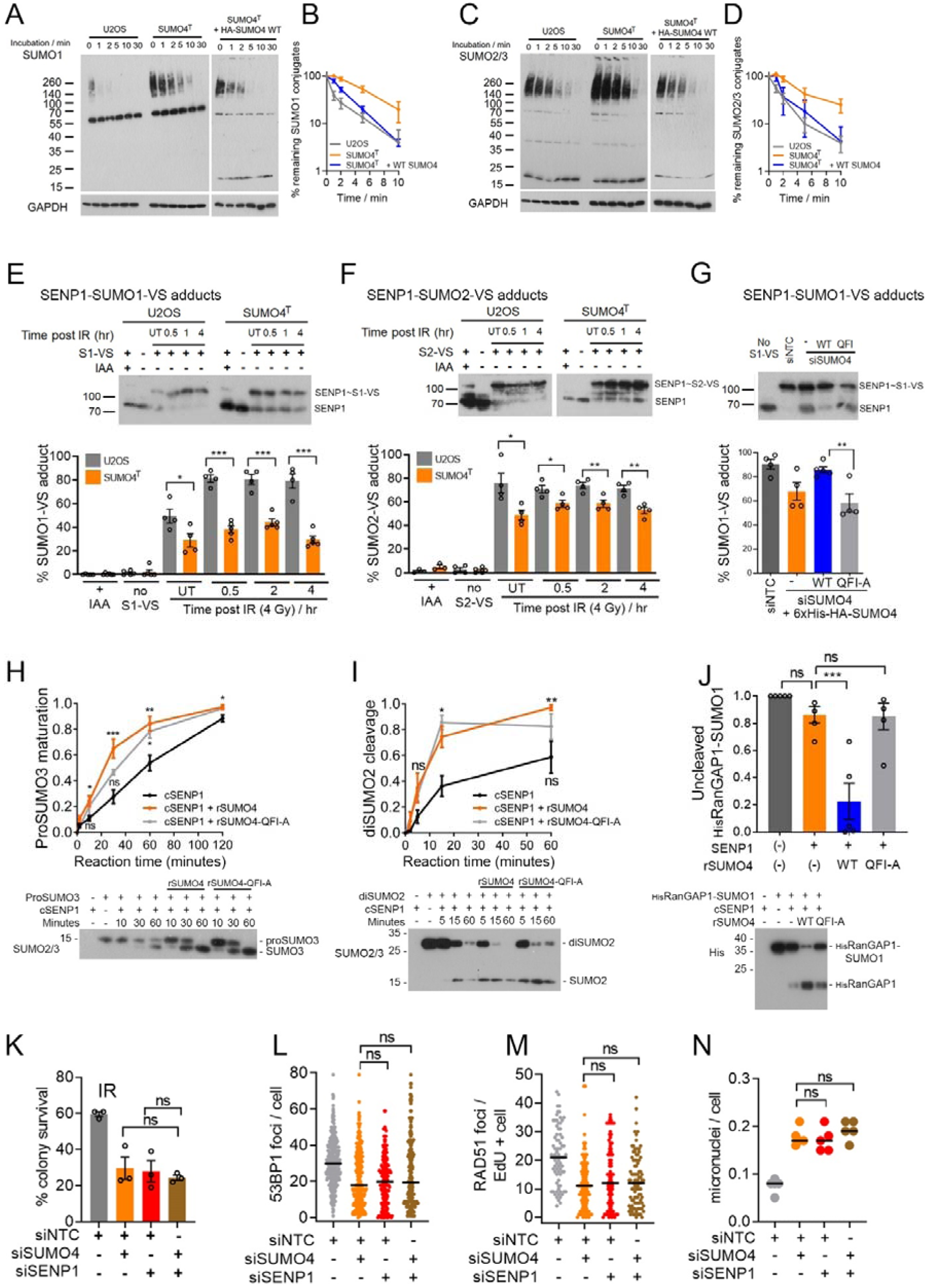
SUMO4 modulates SENP1 SUMO protease activity. **A-D)** Analysis of SUMO-conjugate turnover. Immunoblot of SUMO1 **(A)** or SUMO2/3 **(C)** in U2OS, SUMO4^T^ cl1.11 and SUMO4^T^ cl1.11 expressing, FLAG-HA-SUMO4-WT (induced for 48 hr by doxycycline). At the 0-time point, cells were lysed in a buffer containing the cysteine protease inhibitor, IAA (200 mM), for the preparation of all other time points, they were lysed without IAA. A fraction of the extract without IAA was removed at indicated times and denatured to stop further deconjugation. Lysates were blotted with SUMO1 **(A)** or SUMO2/3 **(C)** antibodies, and the high molecular weight (>70 kDa) signal was calculated by densitometry relative to the 0-time control (lysed in buffer containing IAA and denatured immediately). Graphs show SUMO1 **(B)** or SUMO2/3 **(D)** proteins. Each graph; N=3 experiments, error bars = SEM. **E)** and **F)** HA-SUMO1-vinyl sulfone (S1-VS) **(E)** and HA-SUMO2-vinyl sulfone (S2-VS) **(F),** labelling of SENP1 in U2OS or U2OS SUMO4^T^ cl1.11 cell extracts. Untreated (UT) or after 4 Gy IR at the indicated time points, lysates were incubated with HA-SUMO-vinyl sulfones, then denatured, separated by SDS-PAGE and immunoblotted with SENP1 antibody. The relative amount of upper band (SUMO-VS labelled SENP1) versus unlabelled SENP1 (lower band) was used to calculate % SENP1-SUMO-VS adduct formation. As controls, lysates were incubated with the cysteine protease inhibitor IAA to prevent labelling (+ IAA) or HA-SUMO1-vinyl-sulfone was not included (no S1-VS). Each graph; n= 4 experiments, error bars = SEM. Statistical differences were determined by two-tailed *t*-test. Representative SENP1 blots are shown above each graph. **G)** Quantification of SENP1 HA-SUMO1-vinyl sulfone labelling in U2OS treated with siNTC or siSUMO4 siRNA (48 hr), with the expression of 6xHis-HA-SUMO4-WT or 6xHis-HA-SUMO4-QFI-A induced by doxycycline for 48 hr, n=4, error bars = SEM. A representative immunoblot of SENP1 labelling is shown. Statistical differences were determined by two-tailed *t*-test. **H)** Maturation of Pro-SUMO3 by the SENP1 catalytic domain (SENP1c) in the presence and absence of rSUMO4-WT or QFI-A. 200 nM SENP1 was pre-incubated -/+ 5 µM rSUMO4 at 30°C for 30 minutes prior to the addition of 5 μM Pro-SUMO3 and incubated for the further indicated times before processing and immunoblotting for SUMO2/3. Maturation was calculated as the proportion of the lower SUMO3, cleaved product, of total SUMO3. N=4, error bars = SEM. Statistical significance was assessed by one way ANOVA. A representative immunoblot is shown below. **I)** Cleavage of diSUMO2 by SENP1c in the presence of rSUMO4-WT or QFI-A. 25 nM SENP1c was pre-incubated -/+, 125 nM rSUMO4 for 30 minutes at 30°C before addition of 12.6 μM diSUMO2. The reactions were incubated for the indicated times before processing and immunoblotting for SUMO2/3. Monomeric SUMO2 as a proportion of total SUMO2 was calculated for depolymerization efficiency. N=7, error bars = SEM. Statistical significance was assessed by one way ANOVA. A representative immunoblot is shown below. **J)** DeSUMO-1ylation of RanGAP1-SUMO1 by SENP1c -/+ rSUMO4. 25 nM SENP1c -/+ 1 μM rSUMO4 pre-incubation, then incubated with 1 μM SUMO1ylated His-tagged-RanGAP1(aa398-587) for 5 minutes at 30°C before processing and immunoblotting for His-RanGAP1. The amount of HisRanGAP1-SUMO1 relative to samples without SENP1c are shown. N=5, error bars = SEM. Statistical differences were determined by two-tailed *t*-test. A representative immunoblot is shown below. **K)** Cell survival measured by colony assay of U2OS treated with non-targeting siNTC, siRNA to SENP1 (siSENP1), SUMO4 (siSUMO4) or both for 48 hr before irradiation (2 Gy). N=3, error bars = SEM. Statistical differences were determined by two-tailed *t*-test. **L and M)** Assessment of DNA-damage signalling in U2OS treated with siNTC, siSENP1, siSUMO4 or both for 48 hr before irradiation (4 Gy), fixed 2 hrs later and immunostained for 53BP1 (**L**), or RAD51 (**M**). N=∼150 cells per condition from a total of 3 experimental repeats. Statistical differences were determined by two-tailed *t*-test. **N)** Quantification of micronuclei per cell in U2OS treated with indicated siRNA for 48 hr before irradiation (4 Gy). N=4. Statistical differences were determined by two-tailed *t*-test.

To test whether SUMO4 influences SUMO proteases, we incubated cell lysates with HA-SUMO1- or HA-SUMO2-Vinyl sulfone (SUMO1-VS and SUMO2-VS, respectively). These are active site-directed irreversible inhibitors of SENPs that can act as a proxy for the catalytic activity or catalytic cysteine availability ^55^. We measured the labelling of SENP1, SENP3 and SENP6 SUMO proteases. Of these, we found only SENP1 showed reduced labelling in the SUMO4^T^ lysates (Figures 5E-F and S5A-D). We examined the impact of SENP1 depletion on SUMO2/3 deconjugation, noting a slowing of SUMO2/3 conjugate reduction similar to that observed following SUMO4 siRNA depletion (Figure S5E). Moreover, the expression of SUMO4-WT, but not SUMO4-QFI-A, improved SUMO1-VS labelling of SENP1 in SUMO4-deficient cells (Figure 5G), indicating the presence of SUMO4-WT enhances the accessibility of the SENP1 catalytic site.

These data suggest that SUMO4 positively regulates SENP1 catalytic function. To test if SUMO4 impacts SENP1 protease activity *in vitro*, we generated recombinant SUMO4-WT and QFI-A (rSUMO4) proteins and incubated them with the SENP1 catalytic domain (SENP1c). SENP1c has broad protease activity against multiple SUMO conjugate types ^13^; therefore, we assessed three SENP1 protease activities: proSUMO3 maturation, diSUMO2 cleavage, and deconjugation of SUMO1 from the minimal SUMOylated fragment of RanGAP1 (amino acids 398-587). In each case, incubation of the SENP1c with rSUMO4-WT increased deconjugation activity (Figure 5H-J). rSUMO4-QFI-A was slightly less efficient at promoting SENP1c’s proSUMO3 maturation and diSUMO2 cleavage activity compared to rSUMO4-WT (Figure 5H-J) and the mutant was more profoundly dysfunctional at promoting RANGAP-1 deSUMO1ylation (Figure 5J). The SUMO4 stimulation of protease activity was specific to rSUMO4 as neither rSUMO1 nor rSUMO2 (WT or QFI-A) preincubation increased SENP1c activity against RANGAP1-SUMO1 (Figure S5F). We next made and tested the C-terminal mutants of rSUMO4 (T91X and V94X) and also tested proSUMO2 bearing Proline at codon 90 (i.e. making the C-terminal region of SUMO2 more SUMO4-like). We found rSUMO4 (T91X and V94X) enhanced SENP1c deSUMOylation of RANGAP1-SUMO1, whereas the mutant SUMO2 did not (Figure S5G), suggesting the C terminal ‘P..VY’ residues of SUMO4 are not responsible for promoting SENP1 catalytic activity *in vitro,* and indicating that conjugation ability, in the context of free SUMO, does not suppress SUMO4’s ability to promote SENP1 activity. With the addition of chemical crosslinkers, we were able to detect an association between rSUMO4-WT and SENP1c and a weak association between rSUMO4-WT and rSUMO2 (Figure S5H). Incubation of rSUMO4-WT and SENP1c in the presence of excess SLX4-SIM peptide, but not a peptide in which the hydrophobic residues of the SIM were mutated, reduced the production of the crosslinking rSUMO4-WT∼ complex (Figure S5I), implying that a SIM interface contributes to the formation or stability of the complex.

A role for SENP1 catalytic activity in DSB repair has not previously been defined. To test this, we complemented SENP1 siRNA-treated U2OS with siRNA-resistant forms of SENP1, either SENP1-WT or SENP1-C603A (catalytic active site mutant ^56^). We found that SENP1-WT, but not SENP1-C603A, restored γH2AX, 53BP1 and RAD51 foci numbers to control levels in SENP1 siRNA-treated cells, promoted the survival of U2OS treated with IR, CPT, cisplatin or olaparib and suppressed micronuclei formation (Figure S5J-O). These data suggest that the catalytic function of SENP1 promotes DSB repair. We then assessed the relationship of SENP1 with SUMO4 by performing siRNA co-depletions. In measures of DSB signalling to generate RAD51 and 53BP1 foci, suppression of micronuclei and survival in response to IR, CPT, cisplatin or olaparib, we found that combined SUMO4/SENP1 siRNA treatments were no more deleterious than the impact of each siRNA alone (Figures 5K-N and Figure S5P-R). These data are consistent with the notion that SUMO4 promotes SENP1 catalytic function during DSB repair.

### The SUMO-dependent accumulation of RAP80 at DSBs is regulated by SUMO4-SENP1

Our findings indicate that SUMO4 deficiency disrupts DSB signalling through increasing SUMO conjugates. We hypothesized that a component of the DSB response capable of recognizing SUMO might direct the disruption. The BRCA1-A complex component RAP80 (UIMC1/Ubiquitin Interacting Motif Containing 1) contains tandem SIM-UIM (ubiquitin interaction motif) motifs that interact with SUMO2/3 and K63-ubiquitin chains ^36,37,57^. To determine if RAP80 is affected by SUMO4 disruption, we assessed RAP80 foci in irradiated U2OS cells treated with siRNAs to SUMO1-4. In agreement with prior findings ^36,37^ and the dependence of upstream ubiquitin signalling on SUMO1-3 (Figure 1), depletion of SUMO1 or SUMO2/3 reduced RAP80 foci (Figure 6A). Conversely, SUMO4-siRNA caused a hyper-accumulation of RAP80 foci, which could be reversed by complementation with SUMO4-WT but not by SUMO4-QFI-A (Figure 6B). Similarly, siRNA targeting SENP1, but not siRNAs targeting other SENP proteins, resulted in increased RAP80 foci after IR exposure (Figure S6A). Further, the increased RAP80 foci induced by siRNA SENP1 treatment could be suppressed by SENP1-WT complementation but not by SENP1-C603A (Figures 6C-D), while over-expression of SENP1 reduced IR-induced RAP80 foci numbers in a catalytic-dependent manner (Figure S6B). Co-depletion of SENP1 and SUMO4 did not increase RAP80 foci numbers more than either depletion alone (Figure 6E), consistent with an epistatic relationship between these proteins in promoting DSB repair.

**Figure 6.**
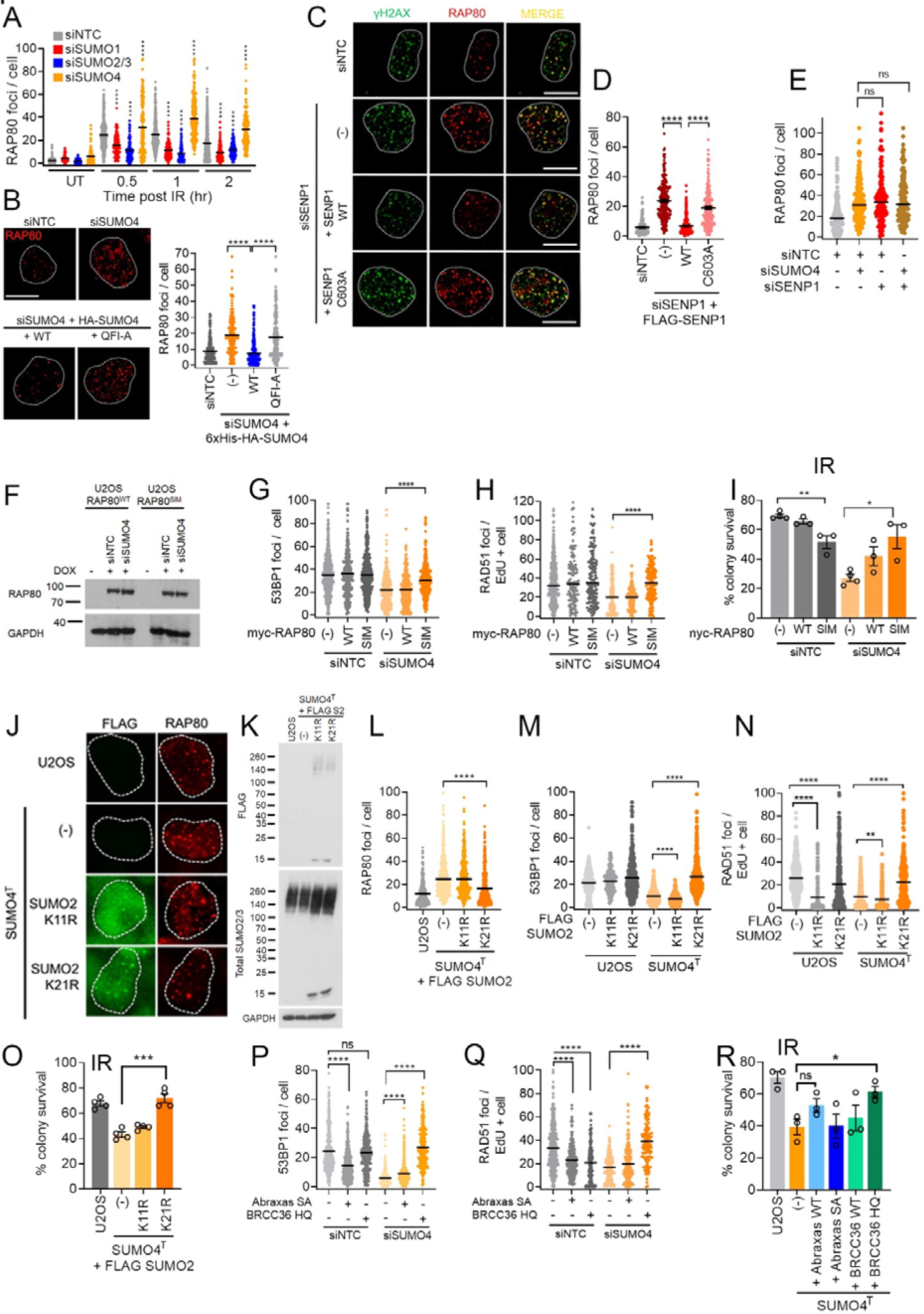
The accumulation of RAP80 is regulated by SUMO4-SENP1. **A)** Quantification of RAP80 foci per cell in U2OS treated with indicated siRNAs for 48 hr before irradiation (4 Gy), fixed at the indicated times and immunostained for RAP80. N >100 cells per condition from three experiments. Statistical difference assessed by two-tailed *t*-test between siNTC and siSUMO for each time point. **B)** Quantification of RAP80 foci in U2OS treated with SUMO4 siRNA and doxycycline to induce expression of 6xHis-HA-SUMO4-WT or 6xHis-HA-SUMO4-QFI-A for 48 hr before irradiation (4 Gy), fixed 2 hrs later. Representative images of cells stained with RAP80 antibody (scale bar = 10 μm) are shown. N ∼150 cells per condition from three experiments. Statistical differences were determined by two-tailed *t*-test. **C and D)** RAP80 foci in U2OS treated with siNTC, or SENP1 siRNA (siSENP1) and doxycycline to induce siRNA-resistant FLAG-SENP1 expression for 48 hr followed by irradiation (4 Gy) and fixed 2 hrs later. **C**) Shows representative images of RAP80 and γH2AX immunostaining, scale bars = 10 μm **D**). Quantification of RAP80 foci per cell from∼150 cells per condition from three experiments. Statistical differences were determined by two-tailed *t*-test. **E)** Quantification of RAP80 foci per cell in U2OS treated with siNTC, siSENP1, siSUMO4 or both for 48 hr before irradiation IR (2 Gy), fixed 2 hrs later and immunostained for RAP80. N=150 cells per condition from a total of three independent experiments. Statistical differences were determined by two-tailed *t*-test. **F)** U2OS lysates from cells bearing doxycycline-inducible myc-RAP80^WT^ and myc-RAP80^SIM^ (SUMO Interacting Motif mutant) treated (+) or not (-) with doxycycline in siNTC or siSUMO4 treated cells (48 hr). The blots were probed for myc and GAPDH loading control. **G and H)** Assessment of 53BP1 (**G**) and RAD51 foci (**H**) in U2OS bearing doxycycline-inducible myc-RAP80^WT^ or myc-RAP80^SIM^ treated with siNTC or siSUMO4 siRNAs for 48 hr, and treated (WT, SIM) or not (-) with doxycycline added 16 hr before irradiation (4 Gy). Cells were fixed 2 hrs later, immunostained for 53BP1 or RAD51, and the number of foci per cell counted. N ∼150 per condition for a total of three experiments. Statistical differences were determined by two-tailed *t*-test. **I)** Cell survival measured by colony assay of U2OS bearing doxycycline-inducible myc-RAP80^WT^ or myc-RAP80^SIM^ treated with NTC or SUMO4 siRNAs for 48 hr, and treated (WT, SIM) or not (-) with doxycycline added 16 hr before irradiation (2 Gy), N=3. Statistical differences were determined by two-tailed *t*-test. **J)** Representative images of RAP80 foci in SUMO4^T^ cl1.11 U2OS expressing doxycycline-inducible FLAG-SUMO2^K11R^ or SUMO2^K21R^ and treated -/+ doxycycline 16 hr before irradiation (4 Gy), fixed 2 hrs later and stained for FLAG (green) or RAP80 (red). **K)** Immunoblot of FLAG-SUMO2 mutants and total SUMO2/3 in SUMO4^T^ cl1.11 cells. **L)** Quantification of RAP80 foci in parental U2OS or SUMO4^T^ cl1.11 expressing doxycycline-inducible FLAG-SUMO2^K11R^ or SUMO2^K21R^ and treated with doxycycline or not (-) before 4 Gy irradiation. For each N ∼150 per condition for a total of three experiments. Statistical differences were determined by two-tailed *t*-test. **M and N)** Quantification of 53BP1 foci (**M**) and RAD51 foci (**N**) in U2OS or U2OS SUMO4^T^ cl1.11 expressing doxycycline-inducible FLAG-SUMO2^K11R^ or SUMO2^K21R^ and treated with doxycycline or not (-) 16 hrs before 4 Gy irradiation. N >100 per condition for a total of three experiments. Statistical differences were determined by two-tailed *t*-test. **O)** Cell survival measured by colony assay following exposure to 2 Gy IR in parental U2OS or SUMO4^T^ cl1.11 cells treated 16 hr with doxycycline to induce expression of FLAG-SUMO2^K11R^ or SUMO2^K21R^ N=4. **P-Q)** Quantification of 53BP1 foci (**P**) and RAD51 foci (**Q**) in U2OS and U2OS SUMO4^T^ cl1.11 treated with NTC or SUMO4 siRNAs for 48 hrs and treated, or not (-) 16 hrs before IR (4 Gy) with doxycycline to induce Abraxas (SA phosphorylation / BRCA1 interaction mutant) or BRCC36 (HQ catalytic inactive mutant) expression. Cells were fixed 2 hrs after IR exposure. N >100 per condition for a total of three experiments. Statistical differences were determined by two-tailed *t*-test. **R)** Cell survival measured by colony assay of irradiated (2 Gy) U2OS SUMO4^T^ cl1.11 treated or not (-) for 16 hr with doxycycline to induce over-expression of Abraxas or BRCC36. Parental U2OS and SUMO4^T^ were used as controls N= 3, error bars = SEM, statistical differences were determined by two-tailed *t*-test.

We assessed the dependence of the accumulations of RAP80 in SUMO4^T^ cells and, as expected, found they were dependent both on upstream K63-Ub signalling components (Figure S6C) and on components of the SUMO conjugation system, including PIAS1 (Figure S6D-E). Further, we tested mutant RAP80, disrupted in its SUMO-interaction motif, F40A/V41A/I42A (hereafter RAP80^SIM^, ^36^) and found that, unlike RAP80^WT^, the SIM mutant did not show increased accumulation in irradiated, SUMO4-depleted cells (Figure S6F). Thus, the increased formation of RAP80 foci, observed following SUMO4 loss, requires active SUMOylation and SUMO-SIM interactions.

We used two approaches to determine if the increased accumulation of RAP80 contributes to the DSB repair defect in SUMO4-deficient cells. Firstly, we tested the over-expression of RAP80^SIM^ since the SIM residues of RAP80 are required to recruit RAP80. Remarkably, RAP80^SIM^ over-expression in SUMO4-deficient cells improved 53BP1 and RAD51 foci accrual and resulted in the restoration of resistance to IR, CPT and olaparib (Figure 6F-I and S6G). Secondly, we tested the impact of SUMO2^K21R^ expression. Biochemical studies using defined lysine SUMO-Ubiquitin linkages suggest K63-ubiquitin dimers conjugated to lysine 21 of SUMO2 are preferentially recognized by the SIM-UIM module of RAP80 ^58^. We found that RAP80 foci numbers in irradiated SUMO4^T^ cells were suppressed to parental levels when SUMO2^K21R^ mutant but not when SUMO2^K11R^ mutant was expressed (Figure 6J-L), consistent with a requirement for SUMO2^K21^ in RAP80 recruitment. Further, SUMO2^K21R^ expression was associated with increased 53BP1 and RAD51 foci after IR and increased resistance to IR, CPT and olaparib of SUMO4-deficient cells (Figure 6M-O and S6H-I). Collectively, these data suggest that increased SUMO-dependent accumulation of RAP80 contributes to the DSB repair defect in SUMO4-deficient cells.

The BRCA1-A complex comprises RAP80, Abraxas/CCDC98, BRCC36, BRCC45, MERIT4, and a proportion of cellular BRCA1:BARD1 heterodimer^59^. The deubiquitinating enzyme, BRCC36, cleaves K63-Ub chains, while phosphorylated Abraxas interacts with the BRCT repeats of BRCA1 and recruits the BRCA1-BARD1 heterodimer ^60–65^. To test if either of these activities contributes to defective DSB repair in SUMO4-deficient cells, we over-expressed a mutant of BRCC36 in which the catalytic residues were substituted, H124Q/H126Q ^66^ (BRCC36-HQ), and a mutant of Abraxas, in which the target residues of phosphorylation were substituted, S404A/S406A ^67^ (Abraxas-SA). We found that BRCC36-HQ, but not Abraxas-SA restored IR-induced 53BP1 and RAD51 foci in irradiated SUMO4-deficient cells (Figures 6P-Q). Further, the expression of BRCC36-HQ, increased the resistance of SUMO4^T^ and SUMO4 siRNA-treated cells to IR, CPT and olaparib (Figure 6R and S6J-K). The Abraxas-SA mutant restored resistance to CPT and Olaparib, but not IR, in SUMO4-deficient cells (Figure 6R and S6K-L). These data indicate that suppressing aspects of the BRCA1-A complex activity can overcome the deleterious impact of SUMO4 loss.

## Discussion

Our analysis confirms distinct roles for SUMO1 and SUMO2/3 in signalling the DNA double-strand break response ^24,27^ and demonstrates an unexpected role for SUMO4. The SUMO1-3 system is well characterized ^68^, but the indication that SUMO4 is unconjugated, combined with a lack of specific SUMO4 detection reagents ^2^, and few unique peptide sequences in mass-spectrometry analysis, has led to SUMO4 being overlooked.

Here we demonstrate that SUMO4, but not SUMO1 or SUMO2, acts to stimulate SENP1 catalytic activity *in vitro*, and that SUMO4 promotes SENP1 catalytic activity in cells. We suggest SUMO4 is unique within the SUMO family and acts in its free state to potentiate deconjugation by SENP1.

The requirement for the SUMO4 SIM-interaction face in promoting SENP1 protease activity against a model-substrate *in vitro* and in supporting DSB signalling in cells is intriguing. Possible mechanisms of SUMO4 action include suppression of product inhibition, to which SENP1 is subject ^56^ and allosteric activation; we observed that excessive SIM availability suppresses the formation of the SUMO4:SENP1 complex *in vitro*, but whether SUMO4 activation occurs through SENP1, or through the substrate/product SUMOs, awaits further biochemical and structural assessment. While our findings correlate the SUMO4-SIM interface with the regulation of SENP1 *in vitro*, we do not discount non-SENP1 or non-SUMO partners as important in the cellular context.

In acting as an unconjugated ubiquitin-like modifier, SUMO4 bears some similarity to UBL5 (ubiquitin-like protein 5), which is not conjugated due to a di-Tyrosine motif in place of the di-Glycine common to other Ubiquitin-like family members. UBL5 instead signals through non-covalent interactions with partner proteins, including the Fanconi Anaemia component FANCI to promote the interaction with FANCD2 in inter-strand cross-link repair ^69^.

We find a crucial role for SUMO4:SENP1 in restricting the SUMO signalling responsible for RAP80 recruitment and, consequently, for promoting DSB repair. RAP80 is part of the BRCA1-A complex, and expressing mutant components of the complex relieves the poor DNA damage signalling and genotoxin sensitives of SUMO4-deficient cells. These findings imply excessive or prolonged BRCA1-A complex activity is deleterious, consistent with its reported ability to restrict ubiquitin-signalling and DNA resection ^60,61,63–66^. Optimal accumulation of RAP80 to DSBs depends on ubiquitin, SUMO and TRAIP binding ^70,71^, the recruiting substrate(s) are presumed to be modified histones or modified, recruited repair proteins ^71–73^. It is feasible that SENP1 locally deSUMOylates concentrated, modified proteins in a manner similar to the activity of SENP6, which deSUMOylates multiple centromeric proteins to support mitosis^74,75^.

Our findings delineate the SUMO E3 ligase: SUMO protease pair, of PIAS1-SENP1, showing they regulate the SUMO conjugates recognized by RAP80. Our data shows this pair are distinct from the previous E3 ligase:SUMO protease pair of PIAS4-SENP2, which are responsible for MDC1-SUMOylation regulation and subsequent clearance from chromatin in G1 cells ^27,32–35^. Thus, cells initiate at least two distinct waves/sprays of SUMOylation/deSUMOylation in the DSB response. The degree to which other SUMO/deSUMOylation events are part of these waves, or are distinct, remains to be seen. We note that the impact of SUMO4 loss on SUMO-conjugate turnover is not restricted to DNA-damage treated cells but also occurs in untreated conditions (e.g. Figure 4A). Thus, we predict that SUMO4 regulation of SENP1 activity is relevant to SENP1-mediated SUMO conjugate homeostasis beyond DNA damage and repair. Dysregulation of the SUMO machinery contributes to tumorigenesis and drug resistance of various cancers, and both SUMO conjugation and deconjugation enzymes are considered drug targets ^54,76,77^. The discovery of a novel component promoting deSUMOylation brings potential new means to target the SUMO pathway.

## Limitations of the study

This work represents the first evidence that SUMO4 has a functional role in DSB repair. One limitation is that we have tested its over-expression in the context of depleted and genetic knock-out cells. Some of the effects observed reveal the non-physiological impacts of manipulating the SUMO system. For example, overexpression of a non-conjugatable form of SUMO2, SUMO2-GA, can suppress the requirement for SUMO4. As SUMO2 cannot activate SENP1 *in vitro*, we speculate that excessive free SUMO2 may relieve some of the deleterious impacts of increased SUMO-conjugates, for example, by competing for the SIM-binding interface of RAP80. Similarly, while over-expression of WT-SUMO4 has no impact on repair outcomes, over-expression of conjugation-proficient SUMO4 is deleterious to DNA repair. We speculate that due to differing charges of several of its surfaces, SUMO4 incorporation into conjugates may suppress vital protein: protein interactions. We have shown endogenous SUMO4 protein and note its expression levels are below that of free SUMO2/3. Previous peptide identification has noted that peptides unique to SUMO4 in human primary tissue and cell lines are at levels far below those for other SUMO proteins^78^ thus, we anticipate SUMO4 is present at levels below that of SENP1. The precise mechanism by which SUMO4 stimulates SENP1 protease function, and the impact SUMO4 has on diverse cellular SENP1 substrates is currently under investigation.

## Declaration of interests

The authors declare no competing interests.

## Author Contributions

AJG designed the study and undertook blots, cell work, SceI assays, imaging and analysis, AJL generated purified recombinant proteins and performed *in vitro* assays, and western blots. KS performed metaphase spreads, RMD performed SceI reporter assays, and YA performed colony assays. HM: performed siRNA validation. AW: performed cell cycle analysis. GR: optimized native SUMO4 immunoprecipitations, and conducted these experiments for subsequent LC-MS/MS analysis. MM conducted in gel LysC digestion, LC-MS/MS and *in vitro* protein-protein cross-linking. ACL: supervised MM and MS data analysis. JRM oversaw the study. AJG and JRM co-wrote the paper. All authors have commented on and edited the manuscript.

## Acknowledgements

Grant funding: Wellcome Trust 206343/Z/17/Z (AJG, MJ, AW), University of Birmingham (AJL, RMD), BBSRC/MIBTP BB/T00746X/1 (YA). CRUK C8820/A19062 and C8820/A28283 (RMD, KS, GR, MM, HM). The U2OS myc-RNF168 cells were generated by Anoop Singh-Chauhan. We thank the Microscopy and Imaging Services at Birmingham University (MISBU) in the Tech Hub facility for microscope support and maintenance. We thank Jeremy Stark (City of Hope, Duarte U.S.A.) for U2OS DR-GFP and NHEJ-EJ5 cells. UbcH9 (human UBC9) plasmid was a gift from Peter Howley (Addgene plasmid # 8651). pET23a-His-hRanGAP1tail, pET28a-His-hAos1 and pET28b-hUba2-His were gifts from Frauke Melchior (Addgene plasmid # 53139, # 53135 and # 53117). We thank the Advanced Mass Spectrometry Facility at the University of Birmingham for assistance with the mass spectrometry measurements. We thank Ron Hay for helpful discussions.

## Resource Availability

All unique/stable reagents generated in this study are available from the Lead Contact with a completed Materials Transfer Agreement. LC-MS/MS data are available via ProteomeXchange with identifier PXD054695.”. Lead contact: Prof Joanna. R. Morris

## Methods

### Cell culture and stable cell lines, and SUMO4^T^cells

All cell lines were cultured in DMEM supplemented with 10% FBS and 1% Penicillin-Streptomycin. FlpIn stable cell lines were generated using U2OS^TrEx-FlpIn^ (a gift from Grant Stewart, University of Birmingham) cells transfected with pcDNA5/FRT/TO-based vectors and the recombinase pOG44 (Invitrogen) using FuGene6 (Promega) at a ratio of 4 μl FuGENE / 1 μg DNA. After 48 hr, cells were grown in hygromycin selection media (150 µg/ml) until colonies formed on plasmid-transfected plates but not controls. Details of all cell lines used in this study can be found in the key resources table. SUMO4 knocTut U2OS^TrEx-FlpIn^ were generated using two different guide RNAs pSpCas9 (BB)-2A PURO (GenScript) plasmids that target the SUMO4 gene at nucleotides (relative to start codon) 150-171 (gRNA #1) and 186-204 (gRNA #4). For each gRNA, three 10 cm^2^ plates of U2OS^TrEx-FlpIn^ were transfected at 5 μg DNA each, using FuGene6. After 48 hr, cells were treated with puromycin (1 μg/mL) to remove un-transfected cells for a further 48 hr. Selected cells were replated at low density on 15 cm^2^ plates to allow clonal growth in DMEM without puromycin. After 10 days, clones were re-seeded and expanded. Clones were screened by PCR using primers that flank the SUMO4 gene using genomic DNA purified with direct PCR buffer (Viagen). Clones that displayed reduced size of SUMO4 PCR product were sequenced by Sanger sequencing to confirm disruption of the SUMO4 locus. Western blot with FLAG was used to confirm the presence of stably integrated 3xFlagCas9. Clones that were positive for 3XFLAG-Cas9 were discarded to reduce the possibility of off-target editing by overexpressed Cas9 nuclease. SUMO4^T^ clone 1.11 was used for the generation of all complemented cell lines as for the parental U2OS^TrEx-FlpIn^ cell line.

**Colony assays, SceI reporter assays and indirect Immunofluorescence** were performed as previously described ^27^.

### Transfection and plasmids

DNA transfections were performed using FuGene6 (Promega) at a ratio of 4 μl FuGENE / 1 μg DNA on 40% confluent cells. Transfection of siRNAs were typically at 10 nM per sequence (or 5 nM per sequence where two are combined). For dual depletions, a total of 10 nM of each siRNA was used to make 20 nM total. Dharmafect-1 was used at a concentration of 1 mL per mL of media. Details of plasmids can be found in supplemental materials. Details of all plasmids and siRNA sequences used in this study can be found in the key resources table

### Vinyl-sulfone labelling, turnover kinetics

Vinyl-sulfone labelling, U2OS cells plated on 10 cm^2^ dishes were treated as indicated for 48 hr before pelleting in ice old PBS and lysis in 1 mL of buffer (150 mM NaCl 10 mM HEPES pH 7.8, 10 mM KCl, 1.5 mM MgCl_2_, 340 mM Sucrose, 10% glycerol 0.2% NP40, protease and phosphatase inhibitor cocktails) followed by sonication and clarification by centrifugation. HA-SUMO-VS (Biotechne) were diluted in PBS and added at a final concentration of 10 ng in a volume of 100 μL for 20 min incubation at room temperature. Reactions were stopped by the addition of 6x Laemmli buffer and boiled. Turnover kinetics, for each condition 2x10 cm^2^ dishes of U2OS (1x10^6^ cell) were plated, siRNA transfected, and doxycycline-treated for 48 hr. Cells were pelleted in PBS control for each condition and were lysed in 1 μL of buffer (250 mM NaCl, 10 mM HEPES pH 7.8, 10 mM KCl, 1.5 mM MgCl_2_, 340 mM Sucrose, 10% glycerol 0.2% NP40, protease and phosphatase inhibitor cocktails) containing 200 mM IAA. After vigorous mixing by pipette for 30 seconds, 150 μL of the sample was added to 50 μL of Laemmli buffer. For turnover, cells were lysed in buffer without IAA, mixed by pipetting and 150 μL samples mixed at indicated times with Laemmli to stop deconjugation, samples were subsequently sonicated and boiled. Details of all antibodies including concentrations used in this study can be found in the supplemental materials table 1 and key resources table

### Metaphase spreads

Cells were plated on 6 well plates 24 hr prior to irradiation at 2 or 3 Gy. The cells were allowed to recover for 24 hr followed by incubation with Karyomax (colcemid) (0.05 μg/ml) for 16 h. Cells were trypsinized and pelleted at 300g for 5 minutes, followed by resuspension in 5 ml of ice-cold 0.56% KCl and incubated at 37°C for 15 min before pelleting at 300g was resuspended and fixed in 5 mL of fixative (ice-cold methanol: glacial acetic acid (3:1)). Fixative was removed, and 15 μl of cell suspension was dropped onto alcohol cleaned slides. Slides were allowed to dry at least 24 hr and then stained with Giemsa solution (Sigma) diluted 1:20 for 20 min. Slide mounting was performed with Eukitt mounting media (Sigma).

### SENP1 *in vitro* protease assays

Details of recombinant protein purification can be found in the supplemental materials and methods section.

### ProSUMO3 maturation assay

200 nM SENP1 (catalytic subunit; R&D systems) and 5 μM recombinant SUMO4 (rSUMO4) were pre-incubated in 50 mM Tris-HCl pH 8, 20 mM NaCl, and 5 mM DTT at 30°C for 30 minutes prior to the addition of 5 μM proSUMO3 in a 20 μl reaction volume. Reactions were terminated by the addition of 20 μl 4xLaemmli buffer and boiling at 95°C for 10 minutes. Samples were analyzed by western blots using SUMO2/3 antibodies.

### DiSUMO2 cleavage assay

25 nM SENP1 and 125 nM rSUMO4 were pre-incubated in the same buffer at 30°C for 30 minutes prior to the addition of 12.6 μM diSUMO2 (R&D systems) in a 20 μl reaction volume. Reactions were terminated by the addition of 20 μl 4xLaemmli buffer and boiling at 95°C for 10 minutes. Samples were analyzed by western blots using SUMO2/3 antibodies.

### HisRanGAP1(aa398-587)-SUMO1 deSUMOylation assay

HisRanGAP1(aa398-587)-SUMO1 was prepared by combining 200 nM SAE1:SAE2, 500 nM Ubc9, 20 μM HisRanGAP1(aa398-587), 20 μM SUMO1, and 5 mM ATP in 20 mM Hepes pH 7.5, 50 mM NaCl, and 5 mM MgCl_2,_ incubated at 37°C overnight before the addition of 2 μM 2-D08 (Ubc9 inhibitor). HisRanGAP1(aa398-587)-SUMO1 deSUMOylation assays involved preincubation of 25 nM SENP1 with 1 μM rSUMO4 for 30 minutes at 30°C in the same buffer and prior to the addition of 1 μM HisRanGAP1(aa398-587)-SUMO1 for a 20 μl reaction mix. Reactions were terminated by adding 20 μl 4x Laemmli buffer and incubation at 95°C for 10 minutes. Samples were analyzed by western blots using polyHis antibody.

**Liquid chromatography-mass spectrometry and Proteomics analysis** is described in the Supplementary Information.

### Statistics

Statistical analysis was by two-tailed Student’s *t*-test unless otherwise stated. *<p0.05, **p<0.01, ***P<0.005 **** P<0.0001. All centre values are given as the mean and all error bars are standard error about the mean (S.E.M.).

Further Materials and Methods can be found in the Supplemental Materials and Methods.

## Notes

### Competing Interest Statement

The authors have declared no competing interest.

### Summary of Updates

Additional evidence of SUMO4 protein and the impact of SUMO4 gene editing on SUMO4 protein.

