## Supplemental Text and Figures for "SUMO4 promotes SUMO deconjugation required for DNA double-strand break repair"

**Supplemental Figures Materials and Methods and Tables.**

**Supplemental Figures.**

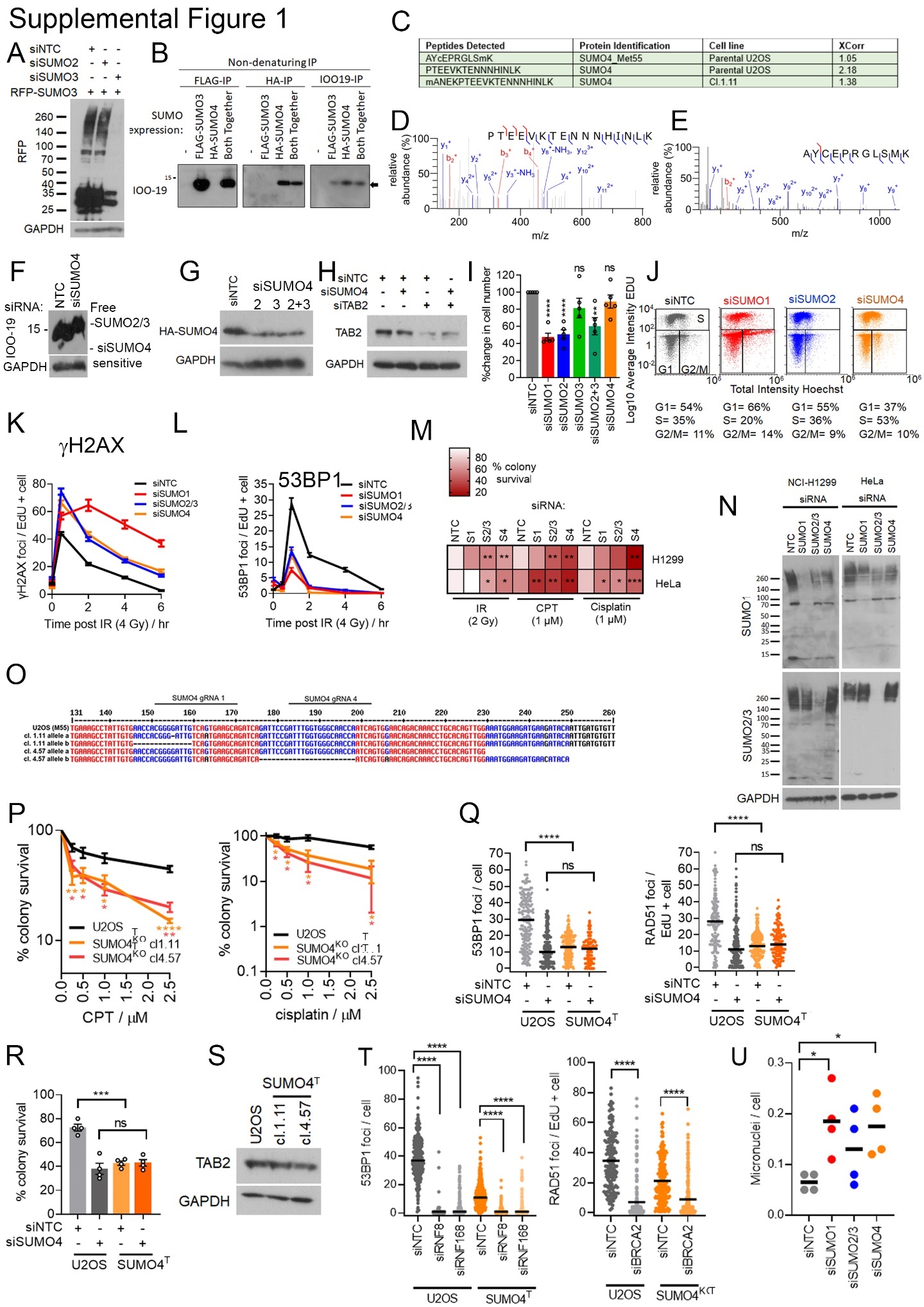

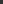

**Supplemental Figure 1. SUMO4 is required for DSB repair.**

**A)** Immunoblot of lysates from U2OS treated with indicated siRNA and RFP-SUMO3 co-transfection for 48 hr prior to lysis.

**B)** Immunoblot of non-denaturing IPs with anti-SUMO2/3/4 antibody IOO-19. Lysates from U2OS cells were transfected with the indicated SUMO3 and/or SUMO4 constructs then split equally and used for FLAG, HA or SUMO4 (IOO-19) IPs. Input lysates and IP elutions were subjected to immunoblotting using the indicated antibodies.

**C)** Table containing peptide sequences detected by LC-MS/MS specific to SUMO-4 from U2OS and SUMO4^T^ cl.1.11 lysates following immunoprecipitation. The XCorr scores for the peptide-spectrum match (PSM) between the MS data and the theoretical spectra are included. Note U20S cells are heterozygous for SUMO4 M55V.

**D)** MS/MS spectra of SUMO4-specific peptide 6-PTEEVKTENNNHINLK-21 from U2OS lysates following immunoprecipitation.

**E)** MS/MS spectra of SUMO4-specific peptide 46-AYcPRGLSMK-56 (whereby c corresponds to a carbamidomethyl modification).

**F)** U2OS treated with indicated siRNA for 48 hr prior to lysis and immunoblotted with the indicated antibodies.

**G)** Immunoblot of lysates from U2OS expressing doxycycline-inducible (1 μg/mL, 48 hr) 6xHis-HA SUMO4 WT (non-siRNA-resistant) cDNA following concomitant SUMO4 siRNA treatment. Lysates were probed with HA and GAPDH (loading control) antibodies

**H)** U2OS treated with indicated siRNA for 48 hr prior to lysis and immunoblotted with the indicated antibodies.

**I)** Number of U2OS cells 72 hr after siRNA treatment for indicated siRNA expressed as a % of siNTC condition. N=4 Error bars = S.E.M, statistical significance by two-tailed t-test.

**J)** Cell cycle profile acquired through high content imaging of U20S 72hr post-siRNA treatment with indicated siRNA. Total Hoechst intensity is plotted against Log10 average Intensity EDU. The gating strategy is indicated by solid black lines. Each representative plot contains data from >15,000 cells from a single biological repeat. The percentage of cells in each cell cycle phase is displayed below each representative graph was obtained from the mean of two biological repeats.

**K)** Number of γH2AX foci / EdU + cell in U2OS treated with indicated siRNA for 48 hr followed by irradiation (4 Gy) and fixation at the indicated time.

**L**) 53BP1 foci counted in EdU+ cells as for **K)**.

**M)** Cell survival measured by colony assay is shown as a heatmap with darker red indicating reduced colony survival in each condition. HeLa and NCI-H1299 cells were treated with indicated SUMO protein siRNAs for 48 hr before treatment with IR (2 Gy), CPT or cisplatin (1 μM 2 hr). Statistical significance is relative to the siNTC for each treatment using two-tailed *t*-test.

**N)** Western blot of indicated siRNA treatments in HeLa or NCI-H1299 cells related to H.

**O)** Sequence alignment of the *SUMO4* gene from genomic DNA recovered from parental U2OS and SUMO4^T^ clones 1.11 and 4.57. Note U2OS are heterozygous for the M55V polymorphism, and the U2OS allele shown here is the WT (Met55) allele. The 1.11 clone (derived using gRNA #1) contains 1 bp deletion and a 14 bp deletion on each allele. The 4.57 clone (derived using gRNA #4) contains a 23 bp deletion and a large deletion that extend beyond the *SUMO4* locus. The 1 bp deletion in clone 1.11 produces sequence alteration from amino acid 49 until a premature stop codon 27 amino acids later. The second allele in clone 1.11 truncates at amino acid 54. Clone 4.57 truncates at amino acids 60 and 76.

**P)** Cell survival of parental U2OS and SUMO4^T^ cl.1.11 (gRNA #1) and cl.4.57 (gRNA #4) clones treated with CPT (left) or cisplatin (right) at the indicated doses for 2 hr before plating for colony growth. N=3-4 * denotes the statistical difference for each dose between parental U2OS and SUMO4^T^ clone, determined by two-tailed t-test

**Q)** U2OS or SUMO4^T^ treated with siNTC or siSUMO4 for 48 hr prior to irradiation (4 Gy) and fixation 2 hr later. Cells were stained for O) 53BP1 or P) RAD51 foci. N >100 cells total from 3 repeats. Error bars = S.E.M, the statistical significance was assessed by students t-test.

**R)** Colony survival in response to IR (2 Gy) in cells treated as for O-P above. Error bars = S.E.M N = 4 and statistical difference by students t-test.

**S)** Western blot of TAB2 and GAPDH in U2OS, and SUMO4^T^ clones 1.11 and 4.57.

**T**) Left: 53BP1 foci in SUMO4^T^ require the canonical upstream DSB repair proteins. U2OS and SUMO4^T^ cl.1.11 were treated with RNF8 or RNF168 siRNA for 48 hr before treatment with IR (4 Gy) and fixation 2 hr later. Cells (~200 total per condition from 3 experiments) were immunostained for 53BP1. Right: RAD51 foci in SUMO4^T^ cells require BRCA2. U2OS and SUMO4^T^ cl.1.11 were treated with siRNA to BRCA2 and stained with RAD51 antibody. The number of RAD51 foci in EdU+ cells (~200 total per condition from 3 experiments) was counted.

**U)** The mean number of micronuclei per cell from 4 experimental repeats in U2OS cells treated with non-targeting siRNA (siNTC) and siRNA to each SUMO protein (siSUMO1-4). N=3 experiments (>50 cells per condition, per experiment).

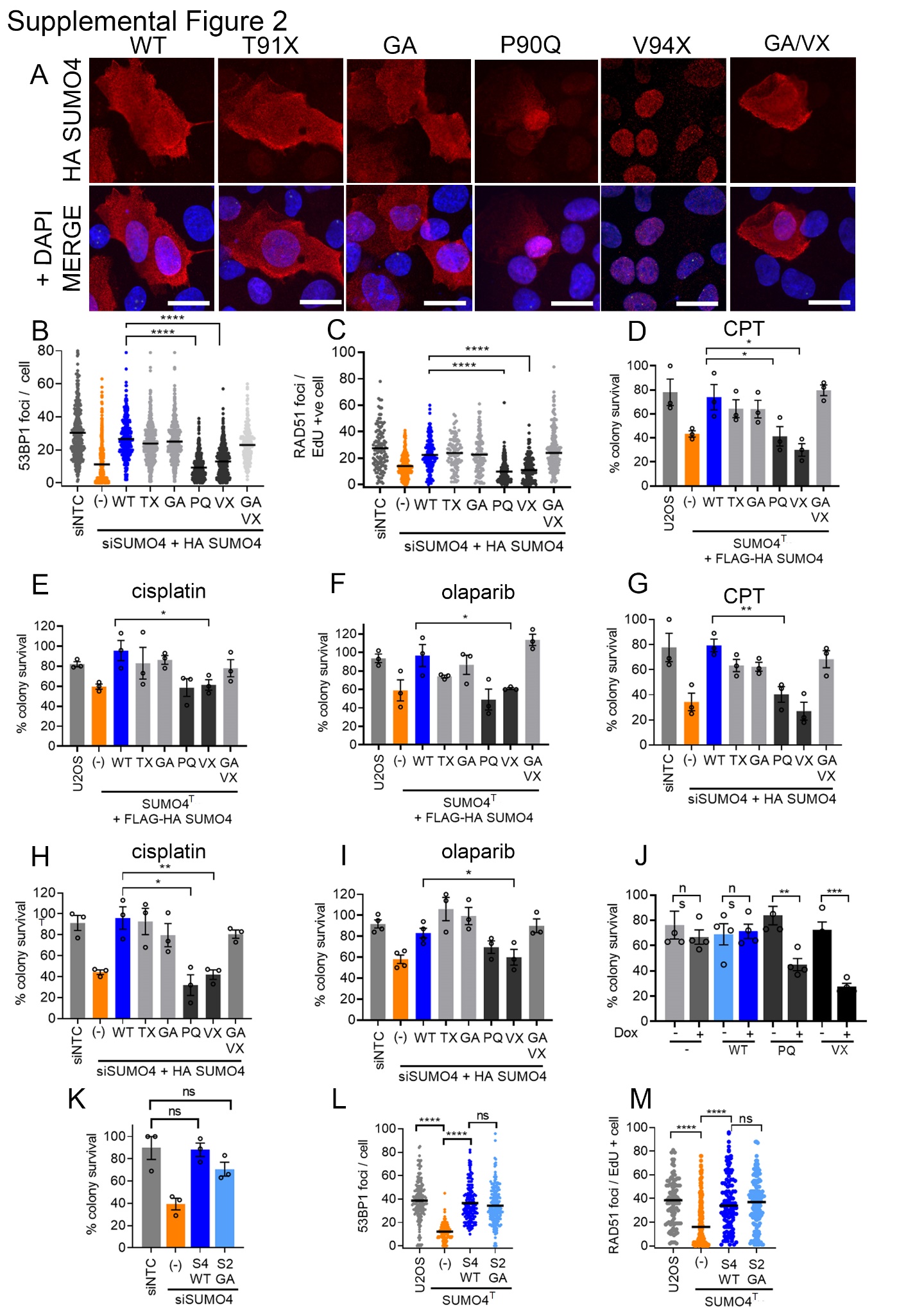

**Supplemental Figure 2. SUMO4 function is suppressed by its conjugation.**

**A)** Indirect immunofluorescence to detect the localization of transiently transfected 6xHis-HA-SUMO4 in U2OS treated with doxycycline to induce expression for 48 hours before fixation and immunostaining with HA antibody. DAPI staining indicates the nucleus. Scale bars = 10 μm.

**B-C)** Impact of 6xHis-HA-SUMO4 C-terminal tail mutants on 53BP1 (**B**) and RAD51 (**C**) foci formation in irradiated cells. U2OS were treated with non-targeting control siRNA (siNTC) or SUMO4 siRNA (siSUMO4) to deplete endogenous SUMO4 with concomitant doxycycline addition to induce expression of siRNA-resistant 6xHis-HA-SUMO4 variants for 48 hr before IR (4 Gy). Cells were fixed 2 hr after IR exposure, followed by immunostaining for 53BP1 or RAD51. N ~150 cells per condition from a total of three experimental repeats.

**D-F)** Assessment of the requirement of the SUMO4 C-terminal tail region on cell survival following exposure to **D)** CPT (1 μM), **E)** cisplatin (1 μM) or **F)** olaparib (10 μM). SUMO4^T^ cl1.11 cells were doxycycline-treated for 48 hr to induce FLAG-HA-SUMO4 protein expression before treatment for 2 hrs with the DNA damaging agents before plating for colony growth. N ~150 cells per condition from a total of three experimental repeats. Statistical differences versus siNTC were determined by two-tailed *t*-test.

**G-I)** as for d-f) but using SUMO4 siRNA and doxycycline to induce expression of siRNA resistant 6xHis-HA-SUMO4 **G)** CPT (1 μM), **H)** cisplatin (1 μM) or **I)** olaparib (10 μM) for 2 hr. N ~150 per condition cells from three experimental repeats. Statistical differences versus siNTC were determined by two-tailed *t*-test.

**J**) U2OS or U2OS stably incorporated for 6xHis-HA-SUMO4-WT, and the SUMO4 mutants 6xHis-HA-SUMO4-P90Q, 6xHis-HA-SUMO4-V94X cells treated with doxycycline for 48 hr to promote SUMO4 over-expression prior to treatment with IR (2 Gy) and plating for colony survival. N=4, error bars = S.E.M and statistical significance by students t-test.

**K**) Colony survival assay of cells treated with 2 Gy of IR prior to plating. Cells were U2OS treated with siNTC or siRNA targeting SUMO4, and untransfected or stably incorporated with FLAG-HA SUMO4-WT or-6xHis-MYC-SUMO2-G92A/G93A (GA) treated with doxycycline for 48 hr to induce the proteins, prior to treatment with IR. N=3 error bars = S.E.M and statistical significance by students t-test.

**L-M)** U2OS, U2OS-SUMO4^T^ stably incorporated for FLAG-HA SUMO4-WT or U2OS-SUMO4^T^ 6xHis-MYC-SUMO2-G92A/G93A (GA) were treated with doxycycline for 48 hr to express the proteins prior to treatment with IR (4 Gy) and fixed 2 hr later. **L**) cells stained for 53BP1 and **M**) RAD51, n= >100 cells from three experiments per condition, error bars = S.E.M and statistical significance by students t-test.

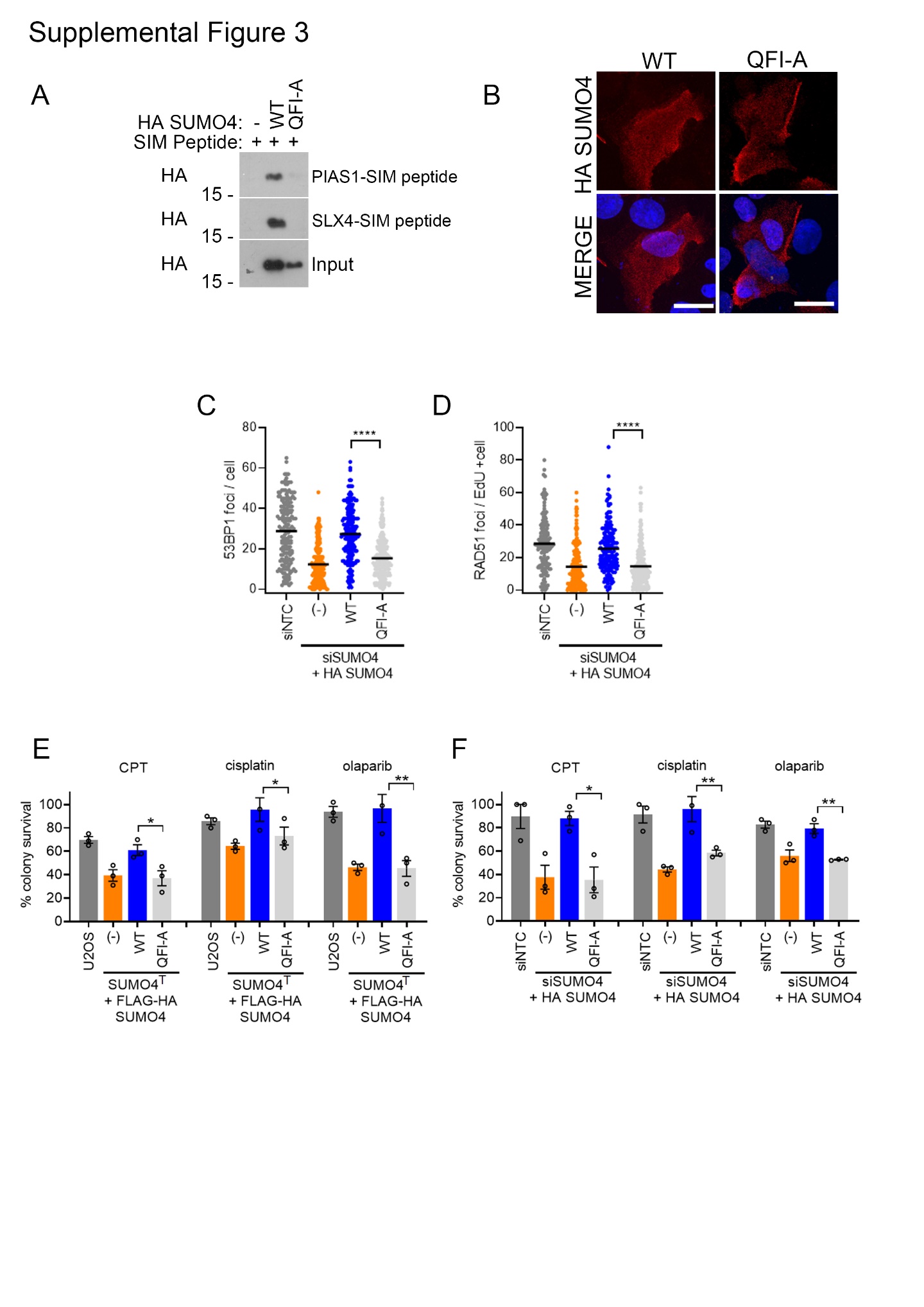

**Supplemental Figure 3. SUMO4 requires its SIM binding groove to promote DSB repair.**

**A)** Western blot analysis of a SIM peptide pull-downs using lysates from U2OS transiently transfected with 6xHis-HA-SUMO4 or 6xHis-HA-SUMO4-QFI-A. U2OS lysates were exposed to Biotin-PIAS1/SLX4-SIM peptides bound to streptavidin beads and western blots were probed with a HA antibody.

**B**) Indirect immunofluorescence of transiently transfected U2OS with 6xHis-HA-SUMO4 variants stained with HA antibody (red). The nucleus is indicated in blue. Scale bars = 10 μm.

**C-D**) Impact of the SIM-binding grove mutant, SUMO4-QFI-A, on 53BP1 (**C**) and RAD51 (**D**) foci formation in irradiated cells. U2OS were treated with non-targeting siRNA (siNTC) or SUMO4 siRNA (siSUMO4) and doxycycline for 48 hr to express 6xHis-HA-SUMO4 variants before exposure to IR (4 Gy) and fixed 2 hrs later. N~150 cells per condition from 3 independent experiments. Statistical differences were determined by two-tailed *t*-test.

**E)** Assessment of the requirement for the SIM-binding face of SUMO4 on cell survival following exposure to CPT (1 μM), cisplatin (1 μM) or olaparib (10 μM) measured by colony assay. U2OS SUMO4^T^ cells were treated with doxycycline for 48 hours to express FLAG-HA SUMO4 variants. Then treated for 2 hrs with the indicated agents and plated for colony formation n=3. Statistical differences were determined by two-tailed *t*-test.

**F)** as for **E)** but using U2OS cells treated with SUMO4 siRNA (siSUMO4) and doxycycline for 48 hr to induce 6xHis-HA-SUMO4 proteins before treatment. Statistical differences were determined by two-tailed *t*-test.

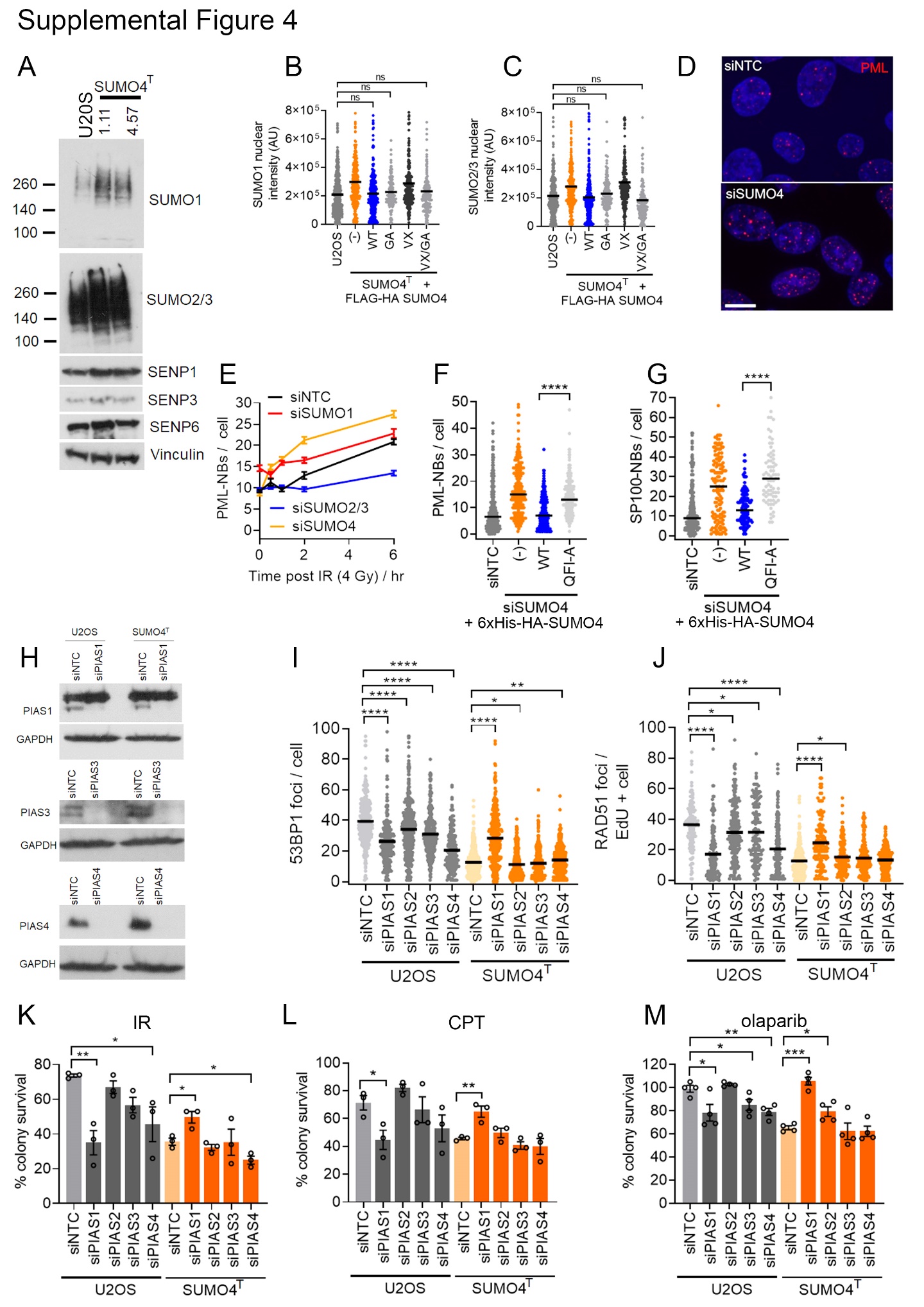

**Supplemental Figure 4. SUMO4 maintains SUMO1-3 homeostasis.**

**A)** Western blot of SUMO1, SUMO2/3, SENP1,3 and 6 in whole cell lysates from two independent SUMO4^T^ clones. Vinculin is included as a loading control.

**B-C**) U2OS, U2OS SUMO4-T cells and U2OS SUMO4-T cells stably incorporating vectors expressing FLAG-HA-SUMO4-WT, or the SUMO4 mutants FLAG-HA-SUMO4-GA, FLAG-HA-SUMO4-VX, and FLAG-HA-SUMO4-VX-GA treated with doxycycline for 48 hr to induce SUMO4 expression followed by permeabilization to remove soluble cytoplasmic and nuclear material and fixed. Cells were stained with SUMO1 antibody (**B**) or SUMO2/3 antibody (**C**) and the intensity of each nucleus was measured on Image J. N = >100 cells, error bars = S.E.M, statistical significance was determined by student t-test.

**D)** Representative images of PML-NBs (red) in U2OS cells treated with non-targeting control siRNA (siNTC) or siRNA to SUMO4 (siSUMO4) 2 hr post 4 Gy IR, counterstained with Hoescht (blue) to reveal the nucleus Scale Bar = 10 μm.

**E)** Quantification of PML-NB numbers in U2OS cells treated with non-targeting control siRNA (siNTC) or siRNA to SUMO1-4 (siSUMO1-4) and treated with irradiation (4 Gy), fixed at the time-points shown and PML bodies counted N=~150 cells per time point from 3 experiments.

**F-G)** PML-NB numbers (**F**) or SP100-NB numbers (**G**) in siNTC or siSUMO4 (48 hr) U2OS concomitantly treated with doxycycline to induce 6xHis-HA-SUMO4 irradiation (4 Gy) and fixed 2 hrs later. N= >75 cells per condition from 3 experiments. Statistical differences were determined by two-tailed *t*-test.

**H)** WB showing siRNA efficiency for the indicated PIAS proteins and GAPDH loading control.

**I-J)** Impact of PIAS siRNAs on 53BP1 (**I**) and RAD51 (**J**) foci formation in irradiated parental and SUMO4^T^ U2OS cells. U2OS and SUMO4^T^ cl1.11 cells were treated with non-targeting control siRNA (siNTC) or siRNA targeting PIAS1-4 (siPIAS1-4) for 48 hr before exposure to IR (4 Gy) and fixed 2 hr later, followed by immunostaining for 53BP1 foci, n =~150 cells per condition from three experiments. Statistical differences were determined by two-tailed *t*-test.

**K-M**) Impact of PIAS siRNA treatment on the survival of U2OS or SUMO4^T^ U2OS cells treated with **K)** IR (2 Gy) or **L)** CPT (1 μM / 2 hr) or **M**) olaparib (10 μM / 2 hr), measured by colony assay. U2OS or SUMO4^T^ cl1.11 cells were treated with non-targeting control siRNA (siNTC) or siRNA to PIAS1-4 (siPIAS1-4) for 48 hrs before treatment with the indicated DNA damaging agents and plating for colony growth. N=3, statistical significances determined by one-way ANOVA.

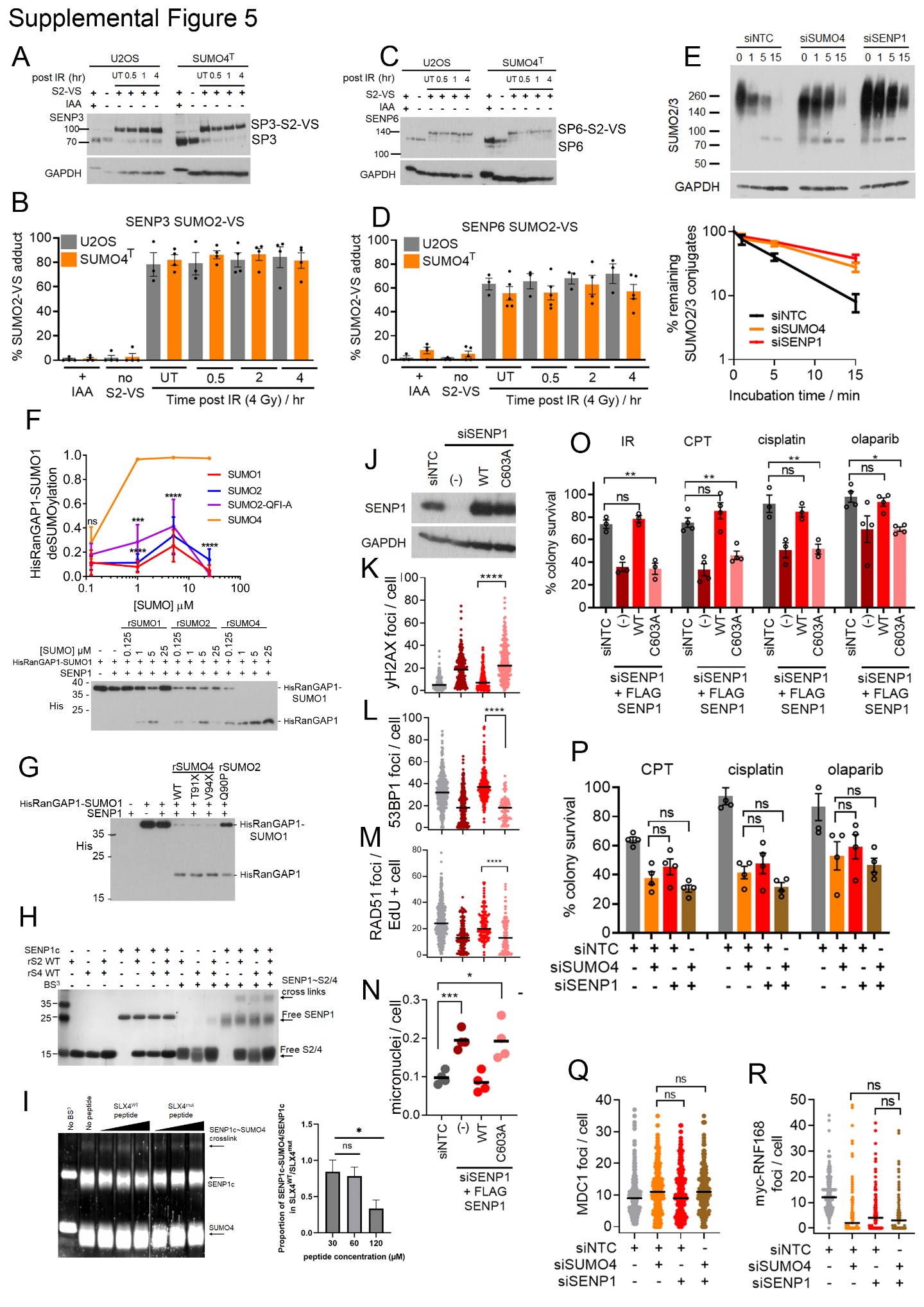

**Supplemental Figure 5. SUMO4 promotes SENP1 protease activity.**

**A)** Representative blot of HA-SUMO2-VS labelling of SENP3 (SP3). HA-SUMO2 vinyl sulfone labelling of SENP3 in cell extracts from U2OS or U2OS SUMO4^T^ that had been either untreated (UT) or irradiated with 4 Gy, and extracts collected at the time-points shown. Reactions were immunoblotted with SENP3 antibody. As controls, lysates were incubated with the cysteine protease inhibitor IAA to prevent labelling (+ IAA) or were not incubated with HA-SUMO2 vinyl-sulfone (no S2-VS).

**B)** The relative amount of upper band (SUMO2-VS labelled SENP3) versus unlabelled SENP3 (lower band) was calculated from 4 experiments. No significant differences between SUMO4^T^ and U2OS lysate SENP3 labelling were found by one-way ANOVA.

**C)** Representative blot of HA-SUMO2-VS labelling of SENP6 (SP6).

**D)** Quantification of the relative amount of upper band (SUMO2-VS labelled SENP6) versus unlabelled SENP6 (lower band) from 4 experiments. No significant differences between SUMO4^T^ and U2OS lysate SENP6 labelling were found by one-way ANOVA.

**E)** Analysis of SUMO2/3 conjugate turnover in U2OS cells treated with non-targeting siRNA (siNTC), siRNA to SENP1 (siSENP1) or to SUMO4 (siSUMO4) for 48 hr. The 1-15 minute time-course lysates were prepared without IAA. A representative immunoblot is shown, (top) and quantification shown (bottom) of high molecular weight (>70 kDa) SUMO2/3 proteins relative to lysates containing IAA at the 0-time point. N=3 experiments, error bars = SEM.

**F)** DeSUMO-1ylation of RanGAP1-SUMO1 by SENP1c -/+ rSUMO1, rSUMO2, rSUMO2-QFIA, or rSUMO4. 0, 0.125, 1, 5, or 25 µM rSUMOs were combined with 25 nM SENP1c at 30°C for 30 minutes, then incubated with 1 μM SUMO1ylated His-tagged-RanGAP1(aa398-587) for 5 minutes at 30°C before processing and immunoblotting for His-RanGAP1. DeSUMOylated HisRanGAP1 as a proportion of total His-RanGAP1 is plotted. N=3, error bars = SEM. Statistical differences were determined by two-way ANOVA. A representative immunoblot is shown below.

**G)** DeSUMO-1ylation of RanGAP1-SUMO1 by SENP1c -/+ rSUMO4 and rSUMO4-C-terminal tail mutants, rSUMO-T91X, rSUMO4-V94X, or ProSUMO2-Q90P. 25 nM SENP1c -/+ 1 μM rSUMO4 were pre-incubated, then incubated with 1 μM SUMO1ylated His-tagged-RanGAP1(aa398-587) for 5 minutes at 30°C before processing and immunoblotting for His-RanGAP1. A representative immunoblot is shown from an N=3.

**H)** SENP1c, SUMO2, SUMO4 proteins were incubated with or without crosslinker bis[sulfosuccinimidyl] suberate (BS^3^) for 30 minutes before quenching with the addition of SDS-PAGE buffer, run on SDS-PAGE and visualised using Coomassie. A representative gel is shown from an n = 2.

**I)** SENP1c, SUMO2, SUMO4 proteins were incubated with or without WT or mutant SLX4 SIM-bearing peptides (between 0.75- and 1.5-times molar excess) and crosslinker bis[sulfosuccinimidyl] suberate (BS^3^) for 30 minutes before quenching with the addition of SDS-PAGE buffer, run on SDS-PAGE and visualised using Coomassie or SYPRO ruby stain (n = 4). A representative SYPRO ruby stained gel is shown (0.5 sec exposure). The graph, right, shows the quantification of the SENP1c~SUMO4 bands. SENP1c~SUMO4 was taken as a proportion of free SENP1c following incubation in the shown quantities of SLX4^WT^ peptide or SLX4^mut^ peptide. Once normalised for loading, bands at the same peptide concentration were compared to indicate the effect of the SLX4 SIM patch on the ­­formation of SENP1c~SUMO4 (where 1 shows no equivalent SENP1c~SUMO4 band as a proportion of free SENP1c when incubated with either peptide, and <1 a reduction in SENP1c~SUMO4 when incubated with WT/mutant peptide). n = 4, error bars = SEM. Statistical significance was determined by two-tailed t-test.

**J)** Immunoblot of SENP1 expression levels in U2OS siRNA-treated with SENP1 siRNA (siSENP1) and complemented with siRNA-resistant FLAG-SENP1 cDNAs for WT or C603A (CA) catalytic mutant SENP1 protein. Anti-SENP1 and anti-GAPDH antibodies were used.

**K-M)** Assessment of the requirement for SENP1 catalytic residues on γH2AX (**K**), 53BP1 (**L**) and RAD51 (**M**) foci in irradiated cells (4 Gy, fixed 2 hr later). U2OS cells with the ability to express siRNA-resistant SENP1 cDNA were simultaneously treated with SENP1 siRNA (siSENP1) and doxycycline to induce SENP1-WT or SENP1-C603A mutant for 48 hrs, before exposure to 4 Gy IR. 2 hrs later the cells were fixed and immunostained with the relevant antibodies and foci counted. n =~150 cells per condition from three experiments. Statistical differences were determined by two-tailed *t*-test.

**N)** Assessment of the requirement for SENP1 catalytic residues on the formation of micronuclei. Cells concomitantly treated, or not, with siRNA to SENP1 and doxycycline to induce siRNA-resistant SENP1 variants, treated with IR (4 Gy). ~100 nuclei per experimental repeat n= 4. Statistical differences were determined by two-tailed *t*-test.

**O)** Requirement for SENP1 catalytic residue C603, to promote cell survival following exposure to several DNA damaging agents as measured by colony assay. U2OS cells with the ability to express siRNA-resistant SENP1 cDNA were simultaneously treated with SENP1 siRNA (siSENP1) and doxycycline to induce SENP1-WT or SENP1-C603A for 48 hrs, before exposure to 2 Gy IR, CPT (1 μM 2 hr), cisplatin (1 μM 2 hr) or olaparib (10 μM 2 hr) before plating for colony survival. N=3, error bars = SEM. Statistical differences were determined by two-tailed *t*-test.

**P)** Assessment of combining SUMO4 and SENP1 siRNA treatment on cell survival in response to CPT (1 μM 2 hr), cisplatin (1 μM 2 hr) or olaparib (10 μM 2 hr). U2OS were treated with non-targeting control siRNA (siNTC) or siRNA to SENP1 (siSENP1), SUMO4 (siSUMO4), or both for 48 hr before exposure to DNA damaging agents and plating for colony growth. N=3, error bars = SEM. Statistical differences were determined by two-tailed *t*-test.

**Q, R)** Assessment of combining SUMO4 and SENP1 siRNA treatment on the formation of MDC1 **(P**) and RNF168 (**F**) foci after irradiation. U2OS cells, or U2OS cells expressing myc-RNF168 (**Q**), were treated with the indicated siRNAs for 48 hr before exposure to 4 Gy IR. Cells were fixed 2 hrs later and immunostained with MDC1 antibody or anti-myc, and foci were counted.

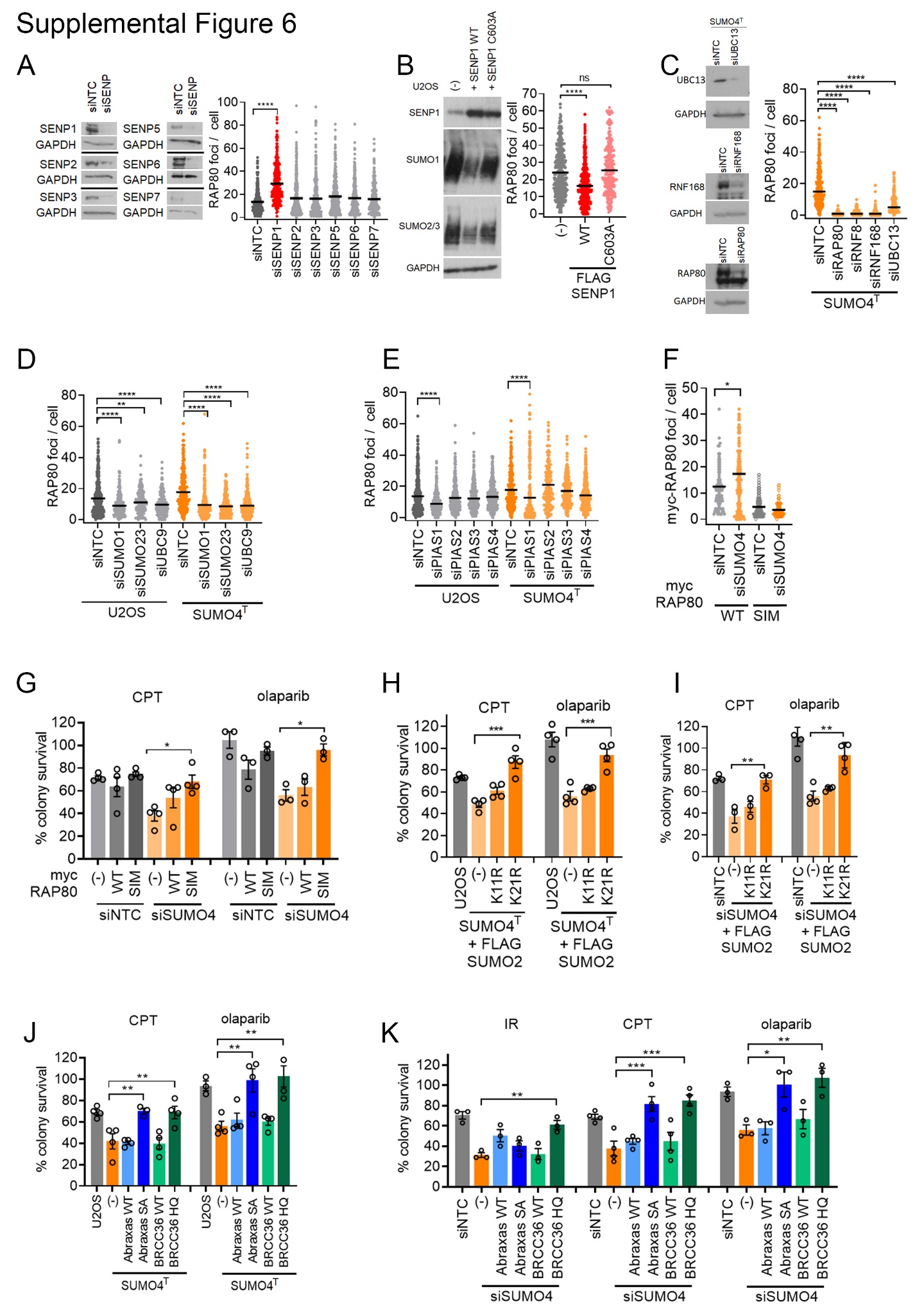

**Supplemental Figure 6. The accumulation of RAP80 at DSBs is regulated by SUMO4:SENP1**

**A)** Assessment of siRNAs to SUMO proteases on RAP80 foci numbers in irradiated U2OS cells. U2OS cells were treated with non-targeting control siRNA (siNTC) or siRNA to each of the SENP-class SUMO proteases (siSENP1,2,3,5,6 & 7) for 48 hr before exposure to IR (4 Gy). They were fixed 2 hrs later, immunostained for RAP80 and the foci counted. N= >100 cells per condition from a total of 3 experiments. Statistical differences were determined by two-tailed *t*-test. Western blots (right) show knockdown efficiency against each SENP in U2OS.

**B)** Requirement for SENP1 catalytic residues to suppress RAP80 foci formation in irradiated cells. Quantification of RAP80 foci (right) in U2OS-SENP1 treated with SENP1 doxycycline (48 hr 1µg / ml) to induce SENP1-WT or the catalytic mutant (C603A), followed by IR (4 Gy). Followed by fixation 2 hr later and stained with antibodies to RAP80. N >100 cells per condition. N=3, Statistical differences were determined by two-tailed *t*-test. Immunoblot (left) of lysates from U2OS-SENP1 treated (+) or not (-) with doxycycline (48 hr 1µg / ml) to induce SENP1-WT or the catalytic mutant (C603A) and probed with antibodies to SENP1, SUMO1, SUMO2/3 and GAPDH.

**C)** Assessment of Ubiquitin signalling in the support RAP80 foci in SUMO4^T^ cells after IR. RAP80 foci (right) in U2OS and U2OS SUMO4^T^ cl1.11 cells were treated with the indicated siRNAs to assess the need for RNF8, RNF168 and UBC13, for 48 hr before irradiation (4 Gy). Cells were fixed 2 hrs later, immunostained with RAP80 antibody and foci counted. N >100 cells per condition from a total of three experimental repeats. Statistical differences were determined by two-tailed *t*-test. Immunoblot (left) shows knockdown efficiency of indicated siRNA in U2OS and U2OS SUMO4^T^ cl1.11.

**D)** Assessment SUMO signalling in the support RAP80 foci in SUMO4^T^ cells after IR. RAP80 foci (right) in U2OS and U2OS SUMO4^T^ cl1.11 cells were treated with the indicated siRNAs to assess the need for SUMO proteins and the SUMO E2 enzyme, UBC9 for 48 hr before irradiation (4 Gy). Cells were fixed 2 hrs later, immunostained with RAP80 antibody and foci counted. N >100 cells per condition from a total of three experimental repeats. Statistical differences were determined by two-tailed *t*-test. Immunoblot (left) shows knockdown efficiency of indicated siRNA in U2OS and U2OS SUMO4^T^ cl1.11.

**E)** siRNA targeting of PAIS1 suppresses RAP80 suppresses excessive RAP80 foci in SUMO4-T cells. U2OS and U2OS SUMO4^T^ cl1.11 cells were treated with the indicated siRNAs for 48 hr before irradiation (4 Gy). Cells were fixed 2 hrs later, immunostained with RAP80 antibody and foci counted. Statistical differences were determined by two-tailed *t*-test. Immunoblot (left) shows knockdown efficiency of indicated siRNA in U2OS and U2OS SUMO4^T^ cl1.11 cells.

**F)** RAP80s SIM is required for its increased accumulation on SUMO4 siRNA depletion. U2OS cells expressing myc-RAP80^WT^ or myc-RAP80^SIM^ were siRNA-treated with non-targeting control (siNTC) or SUMO4 siRNA (siSUMO4) for 48 hrs. Cells were irradiated (4 Gy), fixed 2 hr later and immunostained with myc antibody to detect myc-RAP80 and foci counted n= ~150 cells per condition from three experiments. Statistical differences were determined by two-tailed *t*-test.

**G)** Expression of the RAP80^SIM^ mutant can improve the survival of SUMO4-depleted cells exposed to CPT and olaparib. U2OS cells with the ability to express exogenous RAP80 variants were first treated with non-targeting control (siNTC) or SUMO4 siRNA (siSUMO4) for 48 hrs, followed by 16 hr of doxycycline, or not (-), to induce myc-RAP80^WT^ or myc-RAP80^SIM^ expression before treatment with CPT (1 μM) or olaparib (10 μM) for 2 hr before plating for colony growth. N=3, bars = SEM, and statistical differences were determined by two-tailed *t*-test.

**H)** Expression of the SUMO2^K21R^ mutant can improve the survival of SUMO4^T^ cells exposed to CPT and olaparib. U2OS and SUMO4^T^ cl1.11 cells with the ability to express exogenous SUMO2 variants (SUMO4^T^ FLAG-SUMO2^K11R^ and SUMO4^T^ FLAG-SUMO2^K21R^ cells) were treated or not (-) with doxycycline for 16 hr to induce mutant SUMO2 expression before treatment with CPT (1 μM) or olaparib (10 μM) for 2 hr and plating for colony growth. N=4, bars = SEM, and statistical differences were determined by two-tailed *t*-test.

**I)** Expression of the SUMO2^K21R^ mutant can improve the survival of SUMO4-depleted cells exposed to CPT and olaparib. U2OS cells with the ability to express exogenous SUMO2 variants were first treated with non-targeting control (siNTC) or SUMO4 siRNA (siSUMO4) for 48 hrs, followed by 16 hr of doxycycline, or not (-), to induce FLAG-SUMO2^K11R^ or FLAG-SUMO2^K21R^ before CPT (1 μM) or olaparib (10 μM) treatment for 2 hr and plating for colony growth. N=4, bars = SEM, and statistical differences were determined by two-tailed *t*-test.

**J)** Expression of mutant BRCA1-A complex components can improve the survival of SUMO4^T^ cells exposed to CPT and olaparib. SUMO4^T^ cl1.11 cells engineered for the ability to express exogenous HA-BRCC36 and FLAG-Abraxas variants were treated with dox for 16 hr to induce expression prior to CPT (1 μM) or olaparib (10 μM) treatment for 2 hr and plating for colony growth. Parental U2OS cells were similarly treated. N=3, bars =SEM, and statistical differences were determined by two-tailed *t*-test.

**K)** Expression of mutant BRCA1-A complex components can improve the survival of SUMO4 siRNA treated cells exposed to IR, CPT and olaparib. U2OS cells engineered for the ability to express FLAG-Abraxas, or HA-BRCC36 variants were treated with SUMO4 siRNA (siSUMO4) for 48 hrs followed by doxycycline treatment for 16 hr to induce BRCC36 or Abraxas variant expression before exposure to with IR (2 Gy), CPT (1 μM) or olaparib (10 μM) and plating for colony growth. U2OS cells treated with non-targeting control (siNTC), were also treated with DNA damaging agents N=3, bars =SEM, and statistical differences were determined by two-tailed *t*-test.

**Supplemental Materials and Methods**

**SUMO4 plasmids:** 6xHis-HA-SUMO4 cDNA (NM_001002255.2) was generated by GenScript to include synonymous mutations that render it insensitive to siRNA #2 and siRNA #3. 6xHis-HA-SUMO4 was subcloned into pcDNA5/FRT/TO using HindIII-BamHI sites. 6xHis-HA-SUMO4 non-siRNA resistant vector contains the original cDNA without siRNA resistance. 6xHis-HA-mRFP-SUMO4 were subcloned from the pCDNA5/FRT/TO vector to pcDNA3.1 mRFP using HindIII - XhoI sites. FLAG-HA-SUMO4 constructs were generated using primers that replaced the 6xHis tag with FLAG epitope and were cloned into pCDNA5/FRT/TO using HindIII - XhoI sites.

**SUMO2 plasmids:**6xHis-FLAG SUMO2 K11R and K21R were generated by GenScript and cloned into pCDNA5/FRT/TO at BamHI and XhoI sites.

**SENP1 plasmids:** Human SENP1 cDNA (ENST00000448372.5) was synthesized by GenScript to contain an N terminal FLAG tag and synonymous siRNA resistance mutations to the exon 6 and 12 siRNA used (see table 1). The cDNA also has synonymous mutations to remove BamHI, XhoI and NcoI sites and is cloned into pCDNA5/FRT/TO using BamHI - XhoI sites.

**RAP80, BRCC36 and Abraxas plasmids:** Human RAP80 (also known as UIMC1, ENST00000511320.6) was generated by gene synthesis (GenScript). The cDNA for RAP80 contains an N terminal myc epitope tag. The RAP80 cDNA contains synonymous mutations to make it resistant to siRNA targeting exon 7. Additional synonymous mutations to remove EcoRV, BglII, KpnI, HindIII, and XbaI sites were also introduced. The myc-RAP80 cDNA was cloned into pCDNA5/FRT/TO at BamHI-EcoRV. The SIM binding mutant F40A/V41A/I42A was generated by SDM (GenScript). Human Abraxas and BRCC36 cDNA have synonymous mutations to silence restriction sites and render siRNA resistance and were cloned into pCDNA5/FRT/TO using KpnI - XhoI. Mutations in Abraxas and BRCC36 were introduced by SDM (GenScript).

**SIM-peptide pull-down assay:** U2OS were plated at 1 x 10^6^ cells in 10 cm^2^ plates and transfected with 5 µg pcDNA5 FRT TO-6xHis-HA SUMO4 or pcDNA5 FRT TO-6xHis-HA SUMO4-QFI-A and treated with 4 µg\ml doxycycline for a 48-hour incubation. U2OS cells were lysed in ice-cold RIPA buffer supplemented with protease inhibitors (Roche) and phosphatase inhibitors (Roche). Samples were sonicated at 10 kHz for 2x10 seconds with recovery followed by centrifugation at 14,000g at 4°C for 5 minutes. Magna-bind Streptavidin beads (Thermo Scientific) were washed in TBST and blocked with 20% BSA (TBST) for 2 hours at 4°C with agitation. Each pull-down condition included U2OS lysate, 15 µl streptavidin beads, and 10 µg Biotin-SIM-peptide (PIAS1 or SLX4) subjected to agitation at 4°C overnight. Streptavidin pull-downs were then washed twice in RIPA buffer (protease and phosphatase inhibitors) and then boiled at 95°C in 4xlaemmli buffer. Samples were then centrifuged at 16,000g and analysed using western blots.

**Immunofluorescence:** U2OS were plated at 2.5 x 10^4^ cells/well on 13 mm glass coverslips in 24 well plates (Corning) and attached overnight prior to siRNA depletion for 48 hours. For pre-extraction after 1x PBS wash cells were treated with 250 µL / well ice-cold CSK buffer (100 mM NaCl, 300 mM sucrose, 3 mM MgCl_2_, 0.7% Triton-X100 and 10 mM PIPES) for 30 seconds prior to fixation with 4% Paraformaldehyde (PFA) in PBS at room temperature for 10 minutes. For non-pre-extracted samples, cells were fixed in 4% PFA at room temperature for 10 minutes, followed by permeabilization with 0.5% Triton in PBS for 5 minutes. Coverslips were blocked with 5% FBS in PBS for 1 hour at room temperature, followed by incubation with primary antibodies at 1 µg/mL (or 1:1000 for recombinant and CST MAbs) overnight at 4°C in 5% FBS. Coverslips were washed twice with PBS followed by incubation with Alexa-Fluor 555 conjugated secondary antibodies at 1:2500 for 2 hours at room temperature in the dark. Cells were washed twice with PBS prior to incubation with 250 µL of Hoechst (1 µg/mL) for 2 minutes. Coverslips were mounted on slides using Immuno-Mount (Thermo Scientific) and sealed. Imaging was carried out on a Leica DM6000B microscope using an HBO lamp with a 100 W mercury short arc UV bulb light source. Images were captured at each wavelength sequentially using the Plan Apochromat HCX 100x/1.4 Oil objective at a resolution of 1392x1040 pixels.

**Modified lysis and SDS-PAGE (for SUMO4 separation).** U2OS were seeded in 15 cm^2^ dishes and grown to confluency before scraping into suspension in 1XPBS. Cells were pellet with 500g for 10 minutes and then lysed in 10 mM HEPES-pH 7.6, 200 mM NaCl, 1.5 mM MgCl_2_, 10% glycerol, 0.2 mM EDTA, 1% Triton) supplemented with 1xcOmplete Mini protease inhibitor tablet/2 ml and phosphatase inhibitors, with 4x10 seconds of sonication at 10 kHz with 2 minutes recovery on ice. Lysates were then centrifuged at 13,000g for 5 minutes and the supernatant was combined with 4xLaemmli buffer. Samples were run on a 15% SDS-PAGE gel at 80v for 6-8 hours and transferred to PVDF membrane at 200 mA for 24 hours.

**Purification of recombinant proteins**

**SUMO protein preparation:** BL21(DE3) cells were transformed with pGEX-4T1-GST-ProSUMO3 or pGEX-6P1-GST-SUMO4. Single colonies were used to inoculate 40 ml LB (100 µg/ml ampicillin) and incubated at 37°C. This starter culture was used to inoculate 2-liter LB cultures (100 µg/ml ampicillin) and incubated at 37°C/ 180 rpm to OD595 0.6-0.8. The incubation temperature was reduced to 18°C and 0.5 mM IPTG was added to the cultures for 12-16 hours. BL21(DE3) cells were harvested by centrifugation at 5,000g/ 4°C for 10 minutes. Bacterial pellets were resuspended in ice-cold lysis buffer (20 mM Tris-HCl pH8, 130 mM NaCl, 1mM EDTA, 1% TritonX-100, 10% glycerol, 1 mM DTT, EDTA-free protease inhibitor (Roche)). The bacterial suspension was incubated with 0.5 mg/ml lysozyme for 30 minutes/ 4°C with agitation. 1U/ml DNase (ThermoFisher) was added and samples were sonicated at 20 kHz for 5x30s with 2-minute recovery periods. Samples were centrifuged at 48,000g/ 4°C for 30 minutes. The supernatant was combined with 250 μl Glutathione Sepharose 4B beads (Cytiva) at 4°C for 2 hours with agitation. Samples were centrifuged at 1,000g/4°C/10 minutes and beads were washed thrice with 5 ml lysis buffer and once with 5 ml cleavage buffer. GST-ProSUMO3 purification involved thrombin cleavage buffer (20 mM Tris-HCl pH 8.4, 150 mM NaCl, 1.5 mM CaCl_2_), and GST-SUMO4 purification required PreScission protease cleavage buffer (50 mM Tris-HCl pH 7, 150 mM NaCl, 1 mM EDTA, 1mM DTT) due to a vulnerable thrombin cleavage site unique to SUMO4. GST beads were suspended in 500 μl cleavage buffer and as appropriate 16 U thrombin or 30 U PreScission protease were added for 16 hours/ 4°C with agitation. Samples were centrifuged at 1,000g/ 4°C/ 3 minutes, and the supernatant was isolated for centrifugation at 14,000g/ 4°C for 20 minutes. The supernatant was passed through a 0.45 μm filter before size-exclusion chromatography (SEC) using an AKTA pure™ (UNICORN™ software) Superdex200 Increase 10/300 GL column equilibrated in 20 mM Hepes pH 7.5, 100 mM NaCl, 0.5 mM TCEP: 0.5 ml fractions collected. Eluted fractions corresponding with an increased UV_280_ trace were analysed by SDS-PAGE stained with InstantBlue (Lubioscience). Pure SUMO protein fractions were pooled and stored at -80°C.

**SAE1:SAE2 and RanGAP1(aa398-587) preparation:** BL21 (DE3) were transformed with pET28a-His-hAos1, pET28b-hUba2-His, or pET23a-His-hRanGAP1tail. Single colonies were picked to inoculate 40 ml LB and grown at 37°C. Starter cultures were used to 10 ml starter cultures used to inoculate each litre LB (50 µg/ml kanamycin or 100 µg/ml ampicillin) and grown at 37°C/180 rpm to OD595 ~0.6. Protein overexpression was induced with 1 mM IPTG for SAE1 and SAE2 at 25°C/6 hours and RanGAP1-tail(aa398-587) at 37°C/4 hours. BL21(DE3) cells were harvested by centrifugation at 5,000g/4°C/10 minutes and bacterial pellets were resuspended in 10 ml cold lysis buffer (20 mM Tris/HCl pH 8.0, 300 mM NaCl, 10 mM imidazole, 2 mM β-mercaptoethanol, protease inhibitor). Separately overexpressed SAE1 and SAE2 were combined here. 0.5 mg/ml lysozyme was added and incubated for 30 minutes/4°C/rolling. 1 U/ml DNase was added before sonication at 5x30s at 20 kHz with 2-minute recovery – all on ice. Samples were centrifuged at 48,000xg/4°C/30 minutes and the supernatant was filtered through a 0.45 μm PES membrane (Millex). His-tagged protein lysates were combined with 1 ml nickel beads (Sigma) and incubated at 4°C/2 hours/agitation. Samples were then centrifuged at 1,000g/4°C/10 mins to pellet nickel beads which were resuspended in 10 ml wash buffer (20 mM Tris/HCl pH 8.0, 300 mM NaCl, 50 mM imidazole, 1 mM β-mercaptoethanol, protease inhibitor) ahead of centrifugation as before. Nickel beads resuspended in 5 ml elution buffer (20 mM Tris/HCl pH 8.0, 300 mM NaCl, 500 mM imidazole, 1 mM β-mercaptoethanol, protease inhibitor) and centrifuged as before; supernatant extracted and pushed through 0.45 μm PES filter. This protein suspension was run using an AKTA pure™ (UNICORN™ software) on a HiLoad 16/600 Superdex200 pg column equilibrated with 20 mM Hepes pH 7.5, 100 mM NaCl, 0.5 mM TCEP buffer: 2 ml fractions collected. Fractions corresponding with a UV_280_ peak were analyzed by SDS-PAGE stained with InstantBlue. Fractions containing the purest SAE1:SAE2 or RanGAP1(aa398-587) were pooled and stored at -80°C.

**Ubc9** **purification:** BL21 (DE3) were transformed with pET23a-Ubc9. A single colony was picked to inoculate a 40 ml LB (100 µg/ml ampicillin) started culture, which was grown at 37°C and in turn used to inoculate 2 L LB (100 µg/ml ampicillin). The cultures were grown to an OD595 of 0.6 and Ubc9 overexpression was induced with 1 mM IPTG at 37°C for 4 hours. Cells were harvested by centrifugation at 5,000g/4°C/10 minutes and bacterial pellets were resuspended in 10 ml cold lysis buffer (50 mM Na-phosphate pH 6.5) before lysis using a C3 Emulsiflex. The cell lysate was centrifuged at 48,000g/4°C/30 minutes and the supernatant was filtered through a 0.45 μm PES membrane. The Ubc9 lysate was applied to an SP-sepharose column and latterly washed using the lysis buffer. Ubc9 was eluted from the column using 20 ml Ubc9 elution buffer (50 mM Na-phosphate pH 6.5, 300 mM NaCl, 1 mM DTT, plus cOmplete protease inhibitor) and 1.5 ml fractions were collected. Fractions with the greatest quantity and purity of Ubc9 were pooled and purified by SEC through a Superdex75 equilibrated in transport buffer (20 mM Hepes pH 7.3, 110 mM potassium acetate, 1 mM EGTA, 1 mM DTT, 1 cOmplete protease inhibitor): 4 ml fractions collected. Fractions constituting UV_280_ peak were analyzed by 15% SDS-PAGE and Instantblue stain. Pure Ubc9 protein fractions were pooled and stored at -80°C.

**Chemical cross-linking interaction analysis:** SENP1c (5 µM), SUMO2 (40 µM) and/or SUMO4 (40 µM), were incubated with or without homofunctional NHS-ester crosslinker, bis[sulfosuccinimidyl] suberate (BS^3^) at 800 µM (20 x molar excess to SUMO isoforms). For investigation of the influence of a SIM patch; SENP1c (5 µM), SUMO4 (40 µM), SLX4 peptides WT and mutant peptides at varying concentrations (30, 60 and 120 µM) were all incubated with BS^3^ (800 µM). Peptide sequence information can be found in the KRT. All reactions were conducted in PBS for 30 minutes at room temperature before quenching with the addition of 2 x Laemmli buffer for 15 minutes at room temperature. Samples were boiled at 95°C for 5 minutes and run on 15% SDS-PAGE and visualised using Coomassie stain or SYPRO ruby.

**Cell cycle analysis:** U2OS were plated directly onto 24 well plates (Corning) and attached overnight prior to siRNA depletion for 72 hours. EdU was then pulsed into cells at a concentration of 1 µM per ml for 30 mins before fixation. Cells were directly fixed onto the plate using 4% PFA in PBS for 10 minutes at room temperature, washed in 1x PBS and permeabilized in 0.5% triton for 5 minutes at room temperature. Samples were then blocked in 10% FBS in PBS for 30 mins. Click-iT was performed as detailed in the Click-iT EdU Imaging Kits (Life Technologies). Cells were washed in 1x PBS before incubating with Hoechst for 1 h. Hoechst was removed, and cells were covered with PBS.

Stained plates were imaged on the CellInsight CX5 High-Content Screening (HCS) Platform (ThermoFisher scientific) using the 10x objective and HCS Studio Cell Analysis Software. For each cell the raw values for Total Hoechst intensity and Average EdU intensity were extracted from the software so that it could be plotted.

**Non-denaturing SUMO immunoprecipitation:** 1 x 10^6^ U2OS cells were mock transfected, or transfected with plasmids encoding for FLAG-SUMO3, HA-SUMO4, or both plasmids together. After 72 hours, cells were harvested, washed and lysed in IP Buffer (10 mM HEPES-pH 7.6, 200 mM NaCl, 1.5 mM MgCl_2_, 10% glycerol, 0.2 mM EDTA, 1% Triton) supplemented with protease and phosphatase inhibitors. Following sonication, lysates were cleared (12000 Rpm, 10 minutes, 4°C), antibodies (HA or SUMO2/3/4 IOO-19) or pre-linked agarose beads (FLAG) added and samples rotated (overnight, 4°C). For HA or SUMO4 IPs, protein A/G agarose beads (ThermoFisher Scientific) were added and samples rotated (30 minutes, 4°C). In all cases, beads were then washed 3 times with ice cold IP buffer, before being eluted in 4x SDS Loading buffer and analysed by Western blot. To prepare samples for LC-MS/MS analysis, 1.5 x 10^8^ WT or SUMO4^T^ U2OS cells were subjected to the IP protocol as above.

**Proteomics analysis:** Coomassie strained bands corresponding to the molecular weight of SUMO4 were excised from a Novex^TM^ 4-20% Tris-Glycine Gel (ThermoFisher scientific). Bands were digested as previously described (Aitken and Learmonth, 2002), except 200mM ammonium bicarbonate (pH 8) was used throughout, there was no reduction step, and the bands are digested with 5 ng/μl LysC at 37°C for 16 hours prior to peptide extraction with 100% acetonitrile. The samples were dried down to remove the acetonitrile and then re-suspended in 0.1% formic acid solution in water prior to LC-MS/MS analysis.

Liquid chromatography-mass spectrometry (LC-MS) analysis was performed on an UltiMate® 3000 HPLC series (Dionex, Sunnyvale, CA USA) coupled to a QExactive HF mass spectrometer (ThermoFisher Scientific). The peptides were trapped on precolumn, Thermo Scientific Acclaim PepMap 100 C18 HPLC Columns, 3 µm particle size, 2 cm length, 75 µm I.D., (Dionex, Sunnyvale, CA USA) and separated using a PepMap 100 analytical column (75 µm I.D. x 15 cm, 3 µm) (Dionex, Sunnyvale, CA USA). The LC system was equilibrated in solvent A (0.1% formic acid in water). The gradient used was from 3.2% to 44% solvent B (0.1% formic acid in 100% acetonitrile) for 30 min to elute the peptides, using a flow rate of 350 nLmin^-1.^ All peptides were infused directly into the mass spectrometer via a Triversa Nanomate nanospray source (Advion Biosciences, NY). The capillary voltage was set to 1.7 kV. The mass spectrometer performed a full MS scan (m/z 360−1600) and subsequent HCD MS/MS scans of the 20 most abundant ions with dynamic exclusion setting 15 s. Full scan mass spectra were recorded at a resolution of 120,000 at *m/z* 200 and AGC target of 3×10^6^. For MS/MS, the HCD was set to 28 NCE, Orbitrap resolution to 15,000 and AGC target to 1x10^5^. The width of the precursor isolation window was 1.2 *m/z* and only multiply-charged precursor ions were selected for MS/MS. The MS and MS/MS scans were searched using Protein Discoverer *v.* 2.5 using the Sequest HT algorithm. A fasta file consisting of SUMO1-4 sequences was used (Table 3). The dynamic modifications were oxidation (Met), N-terminal acetylation and N-terminal met-loss+acetylation, with carbamidomethyl (Cys) set as a fixed modification. The number of missed cleavages set to 3. The precursor mass tolerance was set to 10 ppm and the MS/MS mass tolerance 0.02 Da. The PSMs were validated using fixed value PSM validator mode with only high/medium confidence level peptides reported.

**Table 1. Antibodies**

| **Antibody** | **Catalogue number** | **Supplier** | **Concentration / Use** |
| --- | --- | --- | --- |
| FLAG (Gt) | ab1257 | Abcam | 1:1000 (WB) |
| FLAG M2 (Ms) | F1804 | Sigma | 1:2000 (WB, IF) |
| GAPDH 6C5 (Ms) | CB1001 | Calbiochem | 1:5000 (WB) |
| HA.11 (Ms) | 901501 | Biolegend | 1:1000 (WB, IF, IP) |
| PolyHis | H1029 | Merck | 1:5000 (WB) |
| 53BP1 (Gt) | AF1877 | R&D System | 1:2000 (IF) |
| 53BP1 (Rb) | ab36823 | Abcam | 1:2000 (IF) |
| PML C7 (Ms) | ab96051 | Abcam | 1:1000 (IF) |
| PML (Rb) | PA5-79835 | Invitrogen | 1:1000 (IF) |
| SENP1 (Rb) | ab108981 | Abcam | 1:1000 (WB) |
| SENP3 (Rb) | ab124790 | Abcam | 1:1000 (WB) |
| SENP6 (Rb) | HPA024376 | Merck | 1:1000 (WB) |
| SUMO1 Y299 (Rb) | ab32058 | Abcam | 1:1000 (WB, IF) |
| SUMO1 21C7 (Ms) | 33-2411 | Invitrogen | 1:500 (WB, IF) |
| SUMO2/3 8A2 (Ms) | ab81371 | Abcam | 1:1000 (WB, IF) |
| SUMO2/3 12F3 (Ms) | ASM23 | Cytoskeleton | 1:1000 (WB, IF) |
| SUMO4 IOO-19 (Rb) | M06740 | BosterBio | 1:1000 (WB, IP) |
| H2AX-pSer139 (Ms) | ab2893 | Abcam | 1:2000 (IF) |
| H2AX-pSer139 (Rb) | ab22551 | Abcam | 1:2000 (IF) |
| MDC1 (Rb) | PLA-0016 | Bethyl | 1:1000 (IF) |
| MYC (Ms) | ab32 | Abcam | 1:1000 (IF) |
| Vinculin (Rb) | ab129002 | Abcam | 1:2000 (WB) |
| RPA32-pSer33 | ab211877 | Abcam | 1:1000 (IF) |
| RAD51 (Rb) | PC130 | Calbiochem | 1:1000 (IF) |
| SP100 (Rb) | HPA017384 | Atlas | 1:1000 (IF) |
| RAP80 (Rb) | NBP1-87156 | Novus | 1:1000 (IF) |
| BRCC36 (Rb) | 3418-1 | Epitomics | 1:1000 (WB) |
| Ubc9 (Rb) | ab75854 | Abcam | 1:500 (WB) |
| BARD1 (Rb) | ab226854 | Abcam | 1:1000 (IF) |
| RFP (Ms) | 6g6-100 | Chromotek | 1:1000 (WB) |
| Goat α Mouse AF 488 | A11001 | LifeTech | 1:2000 (IF) |
| Goat α Rabbit AF 488 | A11008 | LifeTech | 1:2000 (IF) |
| Goat α Mouse AF 555 | A21422 | LifeTech | 1:2000 (IF) |
| Goat α Rabbit AF 555 | A21428 | LifeTech | 1:2000 (IF) |
| Rabbit α Mouse HRP | P0161 | DAT | 1:10,000 (WB) |
| Swine α Rabbit HRP | P0217 | DAT | 1:10,000 (WB) |

**Table 2 - drugs**

| **Chemical / Treatment** | **Manufacturer / Product code** | **Dosage / Time** |
| --- | --- | --- |
| Camptothecin (CPT) | Merck 208925 | 1 µM 2 hr |
| Doxycycline | Merck D9891 | 1 µg/ mL 72 hr |
| Cisplatin | Selleck S1166 | 1 µM 2 hr |
| Olaparib | Selleck S1060 | 10 µM 2 hr |
| Hygromycin B | Invitrogen H044-81VS | 100 µg/ mL |
| Iodoacetamide (IAA) | Merck I1149 | 200 mM |
| Ionising Radiation | CellRad Irradiator (Precision X Ray) | 2 or 4 Gy |
| ML-792 | Selleck S8697 | 1 µM 3 hr |

**Table S3 –** Protein sequences used in database search for proteomic analysis.

| **Protein** | **Sequence** |
| --- | --- |
| SUMO1 | MSDQEAKPSTEDLGDKKEGEYIKLKVIGQDSSEIHFKVKMTTHLKKLKESYCQRQGVPMNSLRFLFEGQRIADNHTPKELGMEEEDVIEVYQEQTGGHSTV |
| SUMO2 | MADEKPKEGVKTENNDHINLKVAGQDGSVVQFKIKRHTPLSKLMKAYCERQGLSMRQIRFRFDGQPINETDTPAQLEMEDEDTIDVFQQQTGGVY |
| SUMO3 | MSEEKPKEGVKTENDHINLKVAGQDGSVVQFKIKRHTPLSKLMKAYCERQGLSMRQIRFRFDGQPINETDTPAQLEMEDEDTIDVFQQQTGGVPESSLAGHSF |
| SUMO4 – V55 | MANEKPTEEVKTENNNHINLKVAGQDGSVVQFKIKRQTPLSKLMKAYCEPRGLSVKQIRFRFGGQPISGTDKPAQLEMEDEDTIDVFQQPTGGVY |
| SUMO4 – M55 | MANEKPTEEVKTENNNHINLKVAGQDGSVVQFKIKRQTPLSKLMKAYCEPRGLSMKQIRFRFGGQPISGTDKPAQLEMEDEDTIDVFQQPTGGVY |
| SUMO4_cl.1.11A | MANEKPTEEVKTENNNHINLKVAGQDGSVVQFKIKRQTPLSKLMKAYCVSEADQIPIWWATNQWNRQTCTVGNGR |
| SUMO4_cl.1.11B | MANEKPTEEVKTENNNHINLKVAGQDGSVVQFKIKRQTPLSKLMKAYCEPRDCQ |
| SUMO4_cl.4.57A | MANEKPTEEVKTENNNHINLKVAGQDGSVVQFKIKRQTPLSKLMKAYCEPRGLSVKQIRFRFGGQPISGTDKPAQL |
| SUMO4_cl.4.57B | MANEKPTEEVKTENNNHINLKVAGQDGSVVQFKIKRQTPLSKLMKAYCEPRGLSMKQINQ |
